# Tumor invasion as non-equilibrium phase separation

**DOI:** 10.1101/2020.04.28.066845

**Authors:** Wenying Kang, Jacopo Ferruzzi, Catalina-Paula Spatarelu, Yu Long Han, Yasha Sharma, Stephan A. Koehler, Jennifer A. Mitchel, James P. Butler, Darren Roblyer, Muhammad H. Zaman, Jin-Ah Park, Ming Guo, Zi Chen, Adrian F. Pegoraro, Jeffrey J. Fredberg

**Affiliations:** Department of Environmental Science, Harvard T.H. Chan School of Public Health, Boston, MA, USA; Department of Biomedical Engineering, Boston University, Boston, MA, USA; Department of Bioengineering, University of Texas at Dallas, Richardson, Texas; Thayer School of Engineering, Dartmouth College, Hanover, NH, USA; Department of Mechanical Engineering, Massachusetts Institute of Technology, Cambridge, MA, USA; Department of Medicine, Brigham and Women’s Hospital and Harvard Medical School, Boston, MA, USA; Howard Hughes Medical Institute, Boston University, Boston, MA, USA; Department of Physics, University of Ottawa, Ottawa, ON, CA

**Keywords:** breast cancer, invasion, cell migration, jamming, unjamming, phase transition

## Abstract

Tumor invasion depends upon properties of both cells and of the extracellular matrix (ECM). Despite ample evidence that cancer cells can modulate their material state during invasion, underlying biophysical mechanisms remain unclear. Here, we show the potential for coexistence of – and transition between – solid-like, fluid-like, and gas-like phases in invading breast cancer spheroids. Epithelial spheroids are nearly jammed and solid-like in the core but unjam at the periphery to invade as a fluid-like collective. Conversely, post-metastatic spheroids are unjammed and fluid-like in the core and – depending on ECM density – can further unjam and invade as gas-like single cells, or re-jam to invade as a fluid-like collective. A novel jamming phase diagram predicts material phases that are superficially similar to inanimate systems at thermodynamic equilibrium, but here arising in living systems, which exist far from equilibrium. We suggest that non-equilibrium phase separation may provide a unifying physical picture of tumor invasion.

**TWO-SENTENCE SUMMARY:** Using tumor spheroids invading into an engineered three-dimensional matrix, we show here that the cellular collective exhibits coexistent solid-like, fluid-like, and gas-like phases. The spheroid interior develops spatial and temporal heterogeneities in material phase which, depending upon cell type and matrix density, ultimately result in a variety of phase separation patterns at the invasive front, as captured by a jamming phase diagram.

## INTRODUCTION

Invasion of cancer cells from a primary tumor into surrounding tissue is a process wherein migration phenotypes can differ dramatically depending on the properties of the cells as well as those of the surrounding ECM (*1*). It is well established that changes from one migratory phenotype to another depend upon a variety of factors that are cell specific, matrix specific, and interactive (*1, 2*). Current understanding of tumor dynamics begins with a core set of genetic alterations followed by driver mutations, evolution, competition, and resulting sub-clonal heterogeneity within the tumor mass (*3, 4*). Cell invasion and escape from the primary tumor is usually thought to require the epithelial-to-mesenchymal transition (EMT) and associated loss of cell-cell adhesion through loss of E-cadherin. EMT transforms nominally non-migratory epithelial cells into highly migratory mesenchymal cells that can invade individually into surrounding ECM. In multiple models of breast cancer, paradoxically, metastasis requires E-cadherin nevertheless (*5*). A unifying physical picture that describes invasion of cancer cells, either as single cells or as multicellular collectives, is currently lacking.

From tumors of epithelial origin, cells often invade collectively as multicellular strands, sheets, or clusters (*2, 6–8*). Mesenchymal clusters under confinement can also display collective invasion despite their lack of cell-cell adhesions (*9*). As regards the physics of collective invasion, recent evidence implicates the transition of a tumor from a solid-like phase to a fluid-like phase by means of the unjamming transition (UJT) (*10–15*). Experimental work using tumor spheroids suggests that the spheroid core approximates a jammed, solid-like phase in which cellular shapes tend to be rounded and cellular motions tend to be limited, intermittent, and caged by surrounding cells (*11, 12*). The spheroid periphery, by contrast, approaches an unjammed, fluid-like phase in which cell shapes tend to become elongated and cellular motions tend to become larger, more cooperative, more persistent, and sometimes rotational (*16*). Moreover, compared with cells in the spheroid core, cells at the spheroid periphery and invasive branches tend to become systematically softer, larger, longer, and more dynamic (*16*); these mechanical changes arise in part from supracellular fluid flow through gap junctions, suppression of which delays transition to an invasive phenotype (*16*). At the molecular level, increased levels of the small GTPase RAB5A and associated hyper-activation of the kinase ERK1/2 and phosphorylation of the actin nucleator WAVE2 have been implicated in cell unjamming (*11*). Furthermore, the first genome-wide transcriptomic analysis of biological processes that underlie UJT was carried out for the case of human primary bronchial epithelial cells (HBECs) (*17*). Among many other factors, that analysis supports the involvement of cell-ECM adhesions, actomyosin reorganization, activation of ERK and JNK pathways, and downregulation of morphogenetic and developmental pathways (*17*). While that analysis is particular to HBECs, it highlights the fact that the UJT program comprises a coordinated time-dependent interplay of many signaling processes. In HBECs, moreover, UJT is distinct from – and can occur independently of – EMT (*18*). This complex network of interconnected molecular events combines to yield the UJT, which is itself a complex biophysical process resulting from changes to the cell and altered interactions with the surrounding microenvironment.

Using a variety of breast cancer models, Ilina et al. (*15*) recently demonstrated that cell-cell adhesions and mechanical confinement by the surrounding ECM contribute to collective invasion through cell density regulation. They sketched a hypothetical jamming phase diagram in which the interplay between these two factors progressively unjams the tumor from a solid-like non-invasive phase to a fluid-like invading collective, and, finally, to individualized cells that scatter like a gas. Hence, progressive unjamming of a cellular collective by means of the UJT seems to be a major physical route to invasion (*10, 11, 15*). Using multicellular cancer spheroids invading into a three-dimensional (3D) ECM, here we investigate the evolution of distinct material phases within the invasive cellular collective. By quantifying the distribution of cell shapes, packing, and migratory dynamics in both space and time during 3D invasion, we find strong evidence for coexistence of – and transition between – solid-like, fluid-like, and gas-like collective cellular phases, thus confirming earlier findings (*15*). Unlike earlier studies, however, we show further that, depending on cell type and ECM density, routes towards collective invasion can involve not only transitions from jammed to unjammed, but also transitions from unjammed to jammed. A hybrid computational model recapitulates these collective behaviors and results into a novel jamming phase diagram. We propose, further, that this jamming phase diagram may be governed by a small set of ‘effective’ thermodynamic variables, which provide a unifying framework to study the biophysical mechanisms of 3D tumor invasion.

## RESULTS

### In the MCF-10A micro-spheroid, the core approaches a jammed, solid-like phase

#### Structural signatures of unjamming

As a simplified model of an invasive carcinoma, we used MCF-10A epithelial cells (Fig. S1) embedded within an engineered interpenetrating network of Matrigel and Alginate (*16, 19*) (Methods). In these non-malignant cells, ECM stiffness by itself is sufficient to induce a malignant phenotype (*19*). Here we varied the concentration of Alginate without changing the concentration of Matrigel, thus tuning the extracellular environment to a stiffness comparable to malignant breast cancer tissue. Within such matrices, MCF-10A cells spontaneously proliferate, form a micro-spheroid, and, over time, invade into the surrounding ECM (Fig. 1A-B). Previous studies have found that this spontaneous invasive behavior cannot be explained by differences in spheroid size or cell differentiation over time, but rather is attributed to physical changes in cell properties (*16*). We hypothesized, therefore, that the observed collective invasion may be an emergent property associated with cell unjamming. For this reason, we examined evolution of the micro-spheroid at an early stage (days 3-5), when the spheroid is typically 30μm in radius and contains 46 ± 21 cells, and at a later stage (days 7-10), when asymmetric invasive protrusions extend up to 120μm from the spheroid center and the spheroid contains 169 ± 47 cells.

**Fig. 1.**
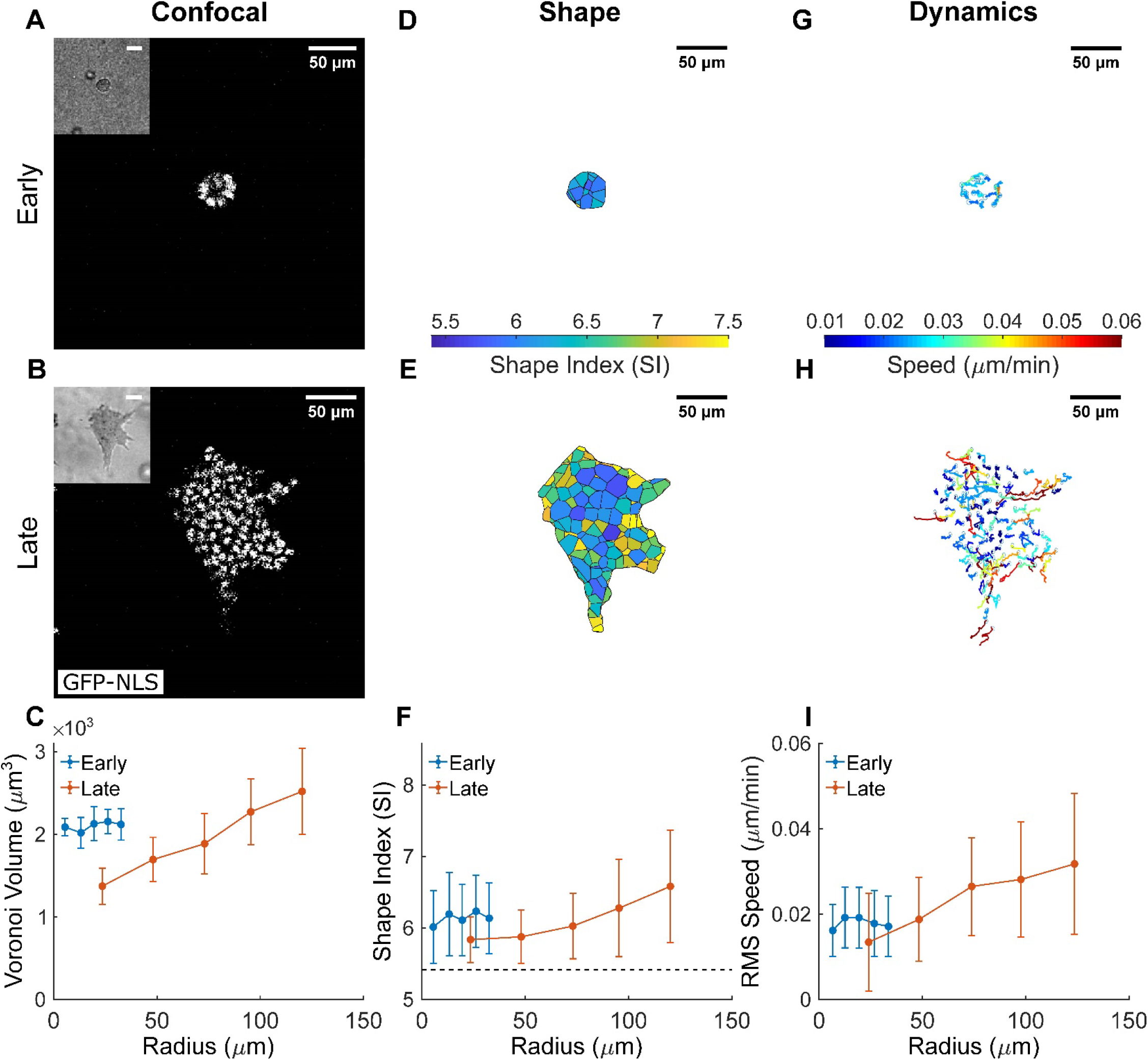
The MCF-10A micro-spheroid exhibits a jammed solid-like core and an unjammed fluid-like periphery. **(A, B)** Equatorial cross-sections of confocal microscopy images show cell nuclei distribution within micro-spheroids grown from GFP-NLS labelled MCF-10A cells at distinct stages of spheroid evolution: early stage (days 3-5; **A**) and late stage (days 7-10; **B**). Micro-spheroids at late stage are much larger compared to the early stage, and show clear invasive protrusions that extend into the ECM. Corresponding cross-sections of bright-field microscopy images outline the micro-spheroid boundary (inset), and are used to generate bounded Voronoi tessellation to estimate cell shape. (**C**) In late-stage spheroids, but not early-stage spheroids, cell volumes obtained by tessellation of nuclear centroids increase with increasing radial position. This result is consistent with previous observations in this model system and attributed to an increase in intra-tumor compressive stress (*16*). (**D, E**) The corresponding cell shapes are shown as 2D cross-sections, color-coded according to their respective 3D Shape Index (SI). Cell SIs exhibit more variability in the late-, than early-stage spheroid. (**F**) Compared to early-stage, cells in the late-stage spheroid core have smaller average SI. In the late-stage spheroid, but not early-stage, SI increased with increasing radial position within the spheroid. This is suggestive of the development of a jammed solid-like core and an unjammed fluid-like periphery. The horizontal dashed line indicates the SI threshold for solid to fluid transition, where proximity away from the threshold (SI > 5.4) suggests transition towards a more fluid-like phase (*24*). (**G, H**) 2D projections of 3D nuclear trajectories tracked over 8 hours reveal that within the early-stage spheroid cell migration is fairly homogenous, whereas in the late-stage spheroid, migratory patterns become highly dynamic. Nuclear trajectories are color-coded according to average migratory speed of the cell over the observation window. (**I**) Compared to the homogenous cell dynamics in the early-stage spheroid, cells in the late-stage spheroid develop a positive radial gradient in migratory dynamics. Consequently, less motile cells are located in the jammed core while more motile cells are located in the unjammed periphery. Data for radial distributions are presented as mean ± STD (n = 5 for both early and late stage spheroids).

To quantify structural characteristics and packing of constituent cells, we identified nuclear locations from 3D confocal microscopy of cells transfected with green fluorescent protein tagged with nuclear localization signal (GFP-NLS). The spheroid boundary was identified from corresponding bright-field images (Fig. 1A-B, insets). Using a bounded Voronoi approach based upon nuclear centroids, we then tessellated the space at each stage of spheroid growth and calculated for each cell the surface area, S, and the volume, V (Methods, Fig. S2). Consistent with previous reports, cell volumes and nuclear volumes varied systematically throughout the spheroid and co-varied linearly (Fig. S3) (*16, 20, 21*). In the late stage micro-spheroid, in particular, we found that cell volumes increased with radial position (Fig. 1C) and cell number densities correspondingly decreased. These gradients have been observed previously and linked to supracellular fluid flows driven by frequently reported gradients of intra-tumor compressive stress (*16*). Compressive stress within tumors is known to become large enough to collapse intra-tumor blood vessels and to increase the metastatic potential of cancer cells (*22, 23*).

We calculated for each cell a non-dimensional shape index (SI), given as the normalized surface area, SI = S/V^2/3^, which can then be used as a structural signature of the degree of local unjamming and fluidization of the collective (*14, 24, 25*), (Methods; Fig. S2). In two-dimensions (2D), cell shapes become progressively more elongated and more variable as the layer becomes progressively more unjammed (*14, 24, 26*). The theory of cellular jamming in 3D suggests that as SI decreases towards a critical value of 5.4 the cellular collective approaches a jammed solid-like phase, while values of SI progressively greater than 5.4 signify collective phases that are progressively less jammed and more fluid-like (*24*). This change is seen to occur in a continuous fashion, akin to jamming or glass transitions, as discussed below, with no discrete, structurally-distinct phase boundary being evident. In the early-stage micro-spheroid, SIs were distributed homogenously with a mean of 6.01 ± 0.51 (Fig. 1D), thereby suggestive of a relatively homogeneous fluid-like phase. In the late-stage micro-spheroid, however, SIs exhibited a clear positive radial gradient (Fig. 1E). Near the spheroid center SI was 5.84 ± 0.32, near the spheroid periphery SI was 6.6 ± 0.79, which is greater than at the center (p<0.01). These regional differences in SI are indicative of the spheroid center being in a material phase in close proximity to a jamming transition, while the spheroid periphery is a material phase that is further removed from a jamming transition (Fig. 1F). At a late stage, therefore, the spheroid center tends to become more jammed and thus solid-like while the periphery tends to become more unjammed and thus fluid-like. We also determined for each cell the 3D principal aspect ratios (AR_1_, AR_2_ and AR_3_), which confirmed the radial trend towards unjamming suggested by the SI (Fig. S4). In all cases the distributions of SI variation from cell-to-cell conformed to a *k-gamma* distribution (Fig. S4). In living systems, inert systems, and computational models, the *k-gamma* distribution has been thought to be yet another structural signature of granular systems approaching a jammed packing (*14, 26, 27*). Using maximum likelihood estimation across all spheroid preparations, the average value of *k* was 10.2 ± 0.1, within the range of values previously reported in inert jammed 3D systems (*27*). We found little variation in *k* between stages of spheroid evolution (Fig. S4). This finding is reminiscent of those reported by Atia et al. in quasi 2D cell layers (*26*), where over a wide range of cell types, *in vivo* and *in vitro*, different underlying pathological conditions, and even different species, *k* was found to fall into a narrow range between 1.9 and 2.5. It remains an open question as to why *k* seems to depend so strongly on dimensionality of the system (*i.e*., 2D versus 3D) but relatively little on the nature of constituent particles.

#### Migratory signatures of unjamming

From image stacks acquired over 8 hours of micro-spheroid growth and cellular migration, we tracked for each cell the nuclear trajectory (Methods). In the early-stage micro-spheroid there was no spatial gradient in cellular migratory speed, whereas in the late-stage micro-spheroid the migratory speed increased systematically from the core to the periphery (Fig. 1I). Cellular motions in the micro-spheroid core were small, sub-diffusive and thus showed evidence of caging (Fig. S5). In contrast, cellular motions in the micro-spheroid periphery were larger, super-diffusive, and thus showed no evidence of caging, consistent with reports from Valencia et al. (*12*). These structural and migratory behaviors further support the interpretation that the center of the more mature spheroid tends to become more jammed and solid-like, whereas the periphery tends to become more unjammed and fluid-like (*11, 12, 16*).

### In the MCF-10A macro-spheroid, the periphery invades as a locally unjammed fluid-like phase

To assess the generality of these results, we examined invasion patterns and cell unjamming signatures in macro-spheroids embedded in matrices spanning a range of collagen densities. Compared to the micro-spheroids described above, these macro-spheroids were larger (extending to a radius of approximately 450μm from the spheroid center) and contained roughly 10-fold to 100-fold as many cells. To form a macro-spheroid we cultured MCF-10A cells on a low attachment substrate in presence of a small volume fraction of Matrigel and allowed the cells to coalesce into a cluster over a period of 48 hours (Methods, Fig. S6). This cluster was then embedded into a self-assembling network of rat-tail collagen I fibrils at either low (2 mg/ml) or high (4 mg/ml) concentration. Using differential interference contrast (DIC) microscopy, these macro-spheroids were imaged continuously as constituent cells proliferated, remodeled the matrix, and initiated invasion. We then used optical clearing (*28*) and multiphoton microscopy (*29*) to obtain stacked images of DAPI-stained nuclei and used second harmonic generation (SHG) signal to obtain images of surrounding collagen (Fig. 2A-B). Within the macro-spheroid, cell-free voids were frequently observed due to the presence of Matrigel (Fig. S6). In these cases, cell shape quantification was restricted to those cells that were either fully surrounded by neighboring cells and/or collagen (Methods, Fig. S2). To allow for comparison between radial distributions of cell shape and migratory dynamics, the latter was assessed using optical flow analysis (*30*) of DIC images (Methods, Movie S1) for the final 8-hour period (40-48 hours).

**Fig. 2.**
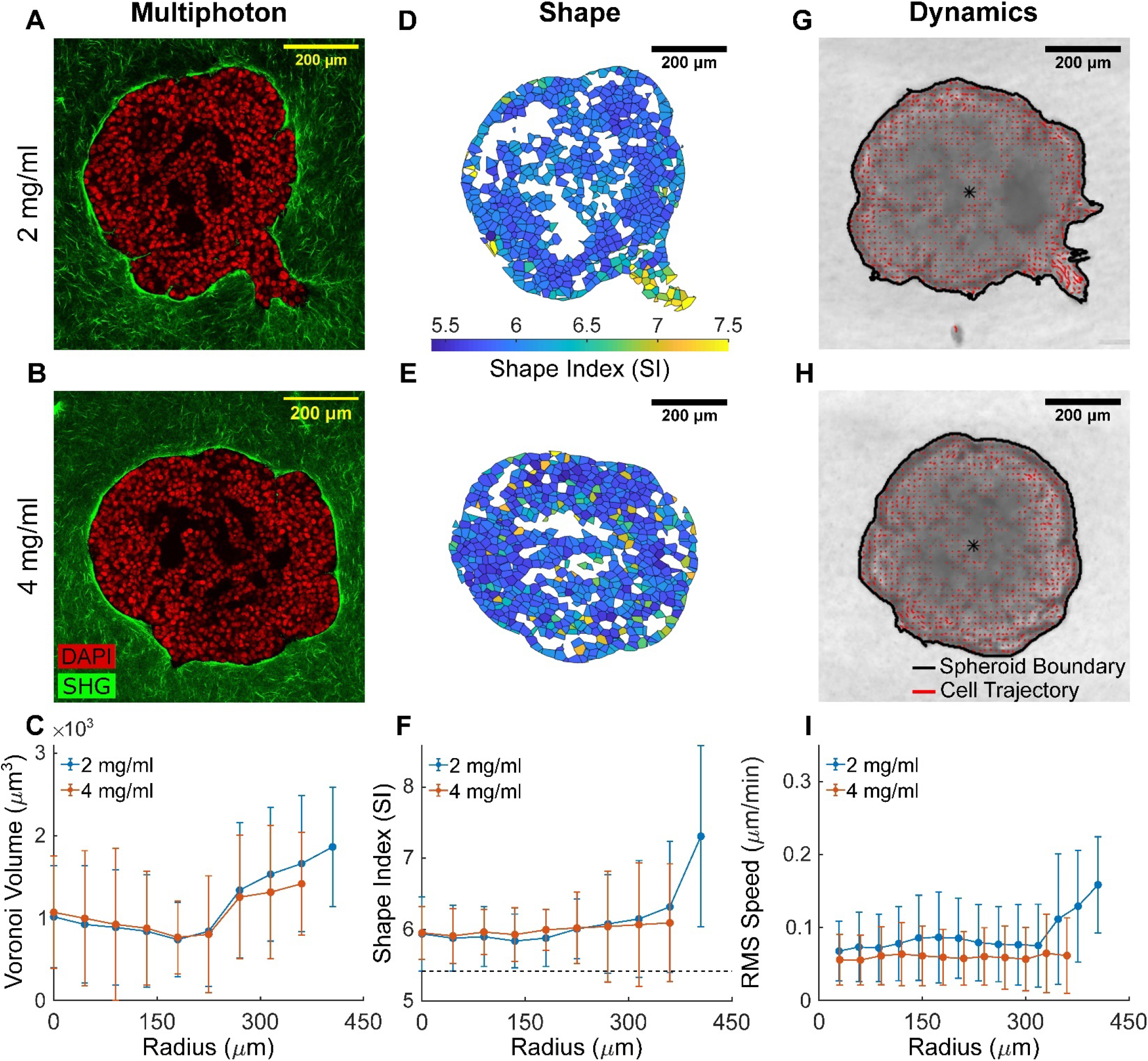
In a manner dependent on collagen concentration, the MCF-10A macro-spheroid locally unjams and fluidizes at the periphery during collective invasion. **(A, B)** Representative equatorial cross-sections of multiphoton images show MCF-10A macro-spheroid behavior when embedded in either 2 (low density) or 4 mg/ml (high density) collagen for 48 hours, with DAPI-stained cell nuclei shown in red and collagen fibers from SHG shown in green. In low density collagen **(A)**, the spheroid develops collective invasive protrusions, while in high density collagen **(B)**, no invasion is observed. Cell-free voids (black) are due to Matrigel used to promote spheroid formation (Methods, Fig. S1); cells immediately neighboring this cell-free region are excluded from subsequent structural analysis (Methods, Fig. S2). (**C**) Similar to observations from the MCF-10A micro-spheroids, Voronoi cell volumes increased from the macro-spheroid core to the periphery. In contrast to the micro-spheroids, average cell volumes from macro-spheroids cultured at both collagen densities are smaller, and suggest that cells in the macro-spheroid experience greater compressive stress (cf. Fig. 1C). (**D-E**) The corresponding cell shapes are shown as 2D cross-sections, color-coded according to their respective 3D Shape Index (SI). Increased and more variable SIs are localized in the region of the spheroid periphery that undergoes collective invasion (**D**). On the other hand, SIs remain narrowly distributed, in the rest of the spheroid periphery and in the core regardless of collagen density. (**F**) In fact, SIs are homogeneously distributed near the threshold for solid-fluid transition (horizontal dashed line indicates solid-fluid transition point at SI=5.4 (*24*)). SIs increased only at the invasive protrusions suggest localized unjamming and fluidization is associated with invasion. (**G-H**) Representative DIC images are shown for MCF-10A macro-spheroids cultured in 2 and 4 mg/ml collagen, with cell migratory trajectories (from optical flow, Methods) superimposed in red. Longer trajectories are observed at the collectively invading regions (**G**). The spheroid boundaries are outlined in black. The entire DIC time-lapse video capturing the dynamics of invasion over 48 hours is shown in Movie S1. (**I**) Radial distributions of average cell migratory speed quantified for the final 8-hour observation window (40-48 hours) conform to expectations from cell shapes. In both collagen densities, migratory speed is homogenously low in the spheroid core, and increased only at sites of localized invasive protrusions. Data for radial distributions are presented as mean ± STD (n = 3 for both 2 and 4 mg/ml spheroids).

At the lower collagen density (2 mg/ml), the MCF-10A macro-spheroids invaded collectively in the form of continuous invasive protrusions and branches (Fig. 2A). By contrast, at the higher collagen density (4 mg/ml), no invasion was observed (Fig. 2B). Much as in the case of the micro-spheroids, in these macro-spheroids cell volumes were smaller near the core compared to the periphery (Fig. 2C). Overall, however, the macro-spheroids exhibited smaller cell volumes than did the micro-spheroids, thereby suggestive of greater compressive stresses in macro-spheroids (*16, 31*). Regardless of changes in collagen density, the spheroid core displayed SIs that were homogenous and small, thus indicating proximity to a solid-like jammed phase. By contrast, in low collagen density the collectively invading regions displayed increased and more variable SIs, indicating localized unjamming and progression towards a more fluid-like unjammed phase (Fig. 2D-F). Migratory dynamics conformed to expectations from cell shapes. After correction for spheroid growth (Methods), migratory speed for cells in low density collagen increased mainly in those regions that had larger SIs and were collectively migrating within invasive branches (Fig. 2G-H). In the epithelial monolayer (*13, 32*) and during 3D spheroid growth (*33*), previous studies have emphasized tissue unjamming and resultant fluidization through the action of cell proliferation. Nevertheless, proliferation seems an unlikely source of the regional unjamming reported here. In our model systems, in fact, changes in collagen density impacted neither spheroid growth nor cell proliferation (Table S1). Instead, we observed larger and more variable cell shapes and faster dynamics restricted to sites where collective invasion occurred.

### In MDA-MB-231 macro-spheroids, the invasive phenotype switches abruptly as a function of ECM density

Localized fluidization of an epithelial cell collective during invasion, such as occurs in the MCF-10A spheroid, involves two main phases: solid- and fluid-like. To account for the possibility that invasion involves also a gas-like phase – corresponding to individually migrating cells after EMT – we performed similar experiments using the post-metastatic cell line, MDA-MB-231. These cells express mesenchymal markers including high Vimentin and low E-Cadherin (*34*) (Fig. S1), exhibit enriched expression in migration-relevant genes (*35*), and form tumors that lead to poor prognosis (*36*), Because spheroid formation in these cells is mediated by integrin β1 adhesion with no cadherin involvement (*37*), these cells form macro-spheroids only in the presence of ECM proteins. Therefore, we formed MDA-MB-231 spheroids by adding 2.5% Matrigel to the cell suspension (*38*) (Methods). For ease of comparison, both MCF-10A and MDA-MB-231 cells were allowed to aggregate using Matrigel, which resulted in macro-spheroids of comparable size (Fig. S6). MDA-MB-231 spheroids were composed of approximately1000-5000 cells and displayed a cell-free core occupied by ECM proteins (Fig. 3; Fig. S6). Restricting our structural analysis to the cells remaining within the continuous tumor mass and fully surrounded by neighboring cells and/or collagen, MDA-MB-231 cells had larger volumes with respect to MCF-10A cells after 48 hours of culture and invasion (Figs. 2C and 3C), larger SIs, (Figs. 2F and 3F) and higher motility regardless of collagen concentration (Figs. 2I and 3I). Therefore, compared to MCF-10A spheroids, cells from post-metastatic MDA-MB-231 spheroids exist in a more motile fluid-like unjammed phase.

**Fig. 3.**
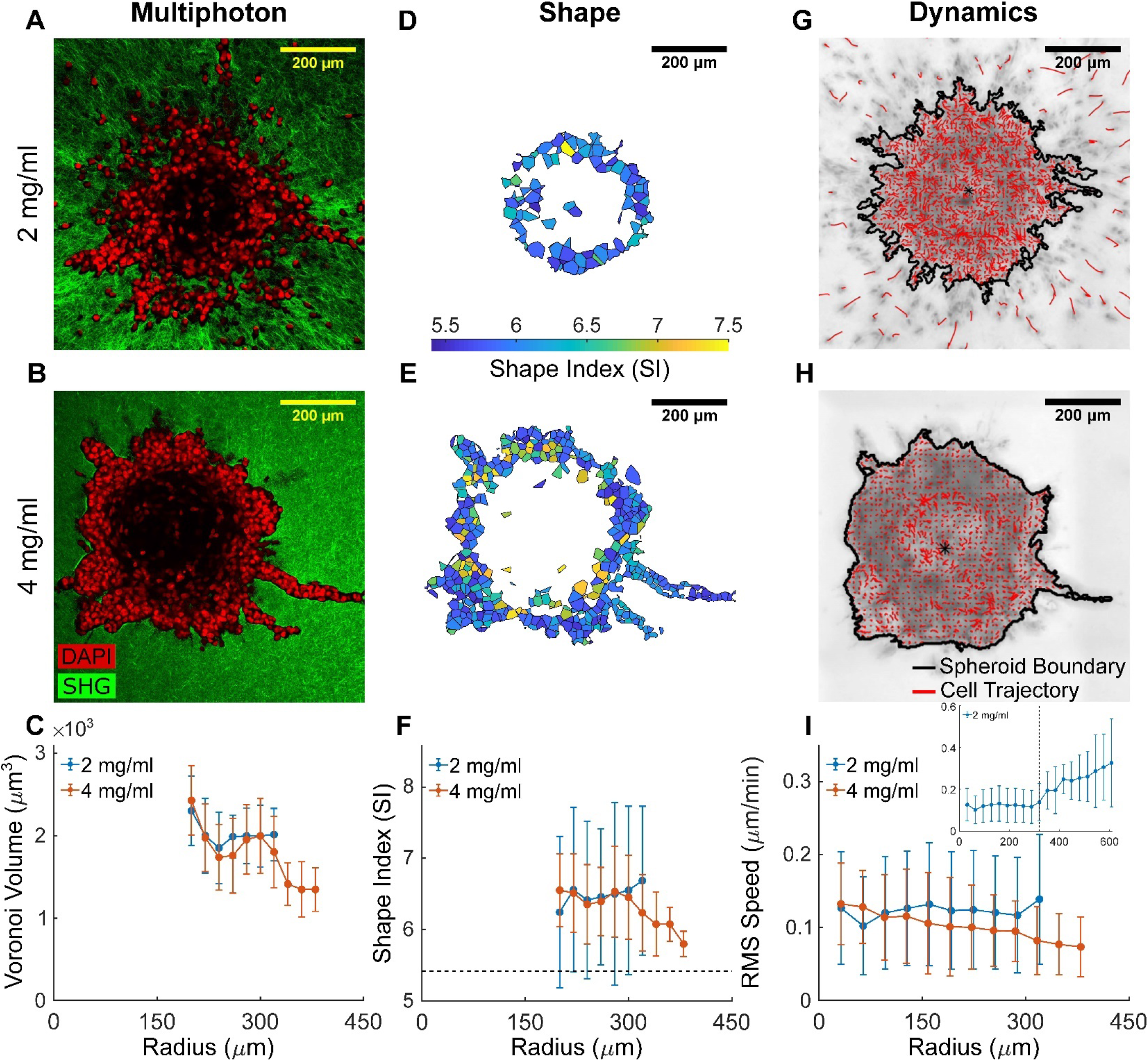
The metastatic MDA-MB-231 spheroid exhibits an unjammed fluid-like phase and undergoes drastically different patterns of invasion depending on collagen concentration. (**A, B**) Representative equatorial cross-sections of multiphoton images show MDA-MB-231 macro-spheroids exhibiting distinct invasion patterns when embedded in low density (2 mg/ml) versus high density (4 mg/ml) collagen for 48 hours. DAPI-stained cell nuclei are shown in red and collagen fibers from SHG are shown in green. In low density collagen **(A)**, these metastatic cells scatter from the spheroid core as individual, gas-like particles. Conversely, in high density collagen **(B)**, single-cell dominant scattering is subdued and invasion is in the form of collective, fluid-like protrusions. We note that the center of MDA-MB-231 spheroids is devoid of cells, as confirmed by staining of histological cross-sections (Fig. S6), and thus result in a hollow shell of highly motile cells rather than a nearly solid spherical structure. Only cells that remain part of the collective are included in the structural analyses (Methods, Fig. S2), hence the absence of data for the first 200 μm of the associated radial distributions. (**C**) Average Voronoi volumes suggest that MDA-MB-231 cells have larger volumes with respect to their MCF-10A counterparts (cf. Fig. 2C). In 2 mg/ml collagen, cell volumes remain roughly independent of radial position. In 4 mg/ml collagen, instead, cell volumes show a decreasing radial gradient. This decrease in cell volume from the spheroid core to the invasive protrusion suggests elevated stress in invading cells from confinement by the collagen matrix. (**D-E**) The corresponding cell shapes are shown as 2D cross-sections, color-coded according to their respective 3D Shape Index (SI). Regardless of collagen concentration, cells from MDA-MB-231 spheroids display higher SI with respect to MCF-10A spheroids (cf. Fig. 2D-E) (**F**) Radial distribution of average SI values is consistent with an unjammed fluid-like phase (horizontal dashed line indicates solid-fluid transition point at SI=5.4 (*24*)). In high density collagen, a radially decreasing gradient in SI suggests that cells jam while invading collectively under matrix confinement. (**G-H**) Representative DIC images are shown for MDA-MB-231 macro-spheroids cultured in 2 and 4 mg/ml collagen, with cell migratory trajectories (from optical flow, Methods) superimposed in red. The spheroid boundaries are outlined in black. The entire DIC time-lapse video capturing the dynamics of invasion over 48 hours is shown in Movie S1. Cell dynamics mirrors structural signatures of cell jamming/unjamming. (**I**) Radial distributions of RMS speed quantified for the last 8-hour observation window (40-48 hours) show that cells in MDA-MB-231 macro-spheroids have homogeneously higher speeds with respect to MCF-10A spheroids (cf. Fig. 2I) and are thus more fluid-like. In low density collagen, cell speed increases further as soon as cells detach from the spheroid and invade as single, gas-like particles (inset, where the radial position of the spheroid boundary is marked by a dashed vertical line). This observation supports the proposed analogy of fluid-to-gas transition. In high density collagen, RMS speed decrease radially with collective invasion, and is supportive of a fluid-to-solid transition due to confinement-induced jamming (*9*). Data for radial distributions are presented as mean ± STD (n = 3 for both 2 and 4 mg/ml spheroids).

Within the MDA-MB-231 macro-spheroid in lower density collagen (2 mg/ml), cell volumes, SIs, and migratory dynamics were nearly homogenous (Fig. 3C, 3F, and 3I). At the spheroid periphery, MDA-MB-231 cells further unjammed by rapidly detaching from the cell collective, reminiscent of evaporating gas particles, and thereafter invaded as single cells along aligned collagen fibers in a highly dynamic manner (Fig. 3I, insert and Movie S1). To our surprise, in higher density collagen (4 mg/ml), cell volumes, SIs, and dynamics displayed a radially decreasing trend (Fig. 3E-F), opposite to that observed in MCF-10A spheroids. Again, migratory dynamics changed in concert with changes in cell shapes (Fig. 3I, Movie S1). While in lower density collagen migratory speed was homogenously distributed within the tumor mass, in higher density collagen one could observe a consistent slow-down of migratory dynamics within the invasive protrusions. We confirmed the presence of such jamming (i.e., a fluid-to-solid) transition by monitoring temporal changes in cell shapes and migratory dynamics in time-lapse experiments of spheroid invasion (Fig. S7). Under conditions of greater confinement by the ECM, as the branch invaded into the matrix both SIs and cell speeds decreased. Overall, we found that – depending on collagen concentration – liquid-like MDA-MB-231 spheroids invade while undergoing liquid-to-gas unjamming (in 2 mg/ml) or a liquid-to-solid jamming (in 4 mg/ml), respectively representing single and collective modes of cell migration.

But how does the transition between these different modes of invasion occur as a function of ECM confinement? Using graded concentrations of collagen (1 to 4 mg/ml), we tracked over time the number of single MDA-MB-231 cells that had detached from the continuous primary spheroid mass, escaped that spheroid, and invaded in a gas-like fashion into the ECM. The number of such single invading cells was found to be not only time-dependent but, more importantly, dependent on collagen concentration (Fig. 4A-C). On day 0 no cell escape was evident at any collagen concentration; immediately after embedding in collagen, all cells remained within the spheroid. On day 1 a modest level of cell escape became evident at lower collagen concentrations (1 and 2 mg/ml) but not at higher concentrations. On day 2 the number of detached invading cells became much larger and highly sensitive to collagen concentration. By day 3, remarkably, the number of detached invading cells stabilized into a striking biphasic switch-like dependence on collagen concentration. The collagen concentration demarking this step-like transition for MDA-MB-231 spheroids fell between 2 and 3 mg/ml. To uncover structural and mechanical features of ECM that might underlie the observed changes in invasive phenotype, we conducted high-resolution multiphoton microscopy imaging and confined compression testing at these graded collagen concentrations (Fig. 4D-G). As collagen concentration increased, bulk compressive energy storage (quantifying the nonlinear material properties commonly referred to as stiffness, cf. Supplementary Methods) did not change whereas fiber density increased and both porosity and hydraulic permeability decreased (Fig. 4D-G; Fig. S8). Thus, our biomechanical characterization suggests that, at least in this experimental system, ECM density can control the mode of invasion independently from ECM stiffness.

**Fig. 4.**
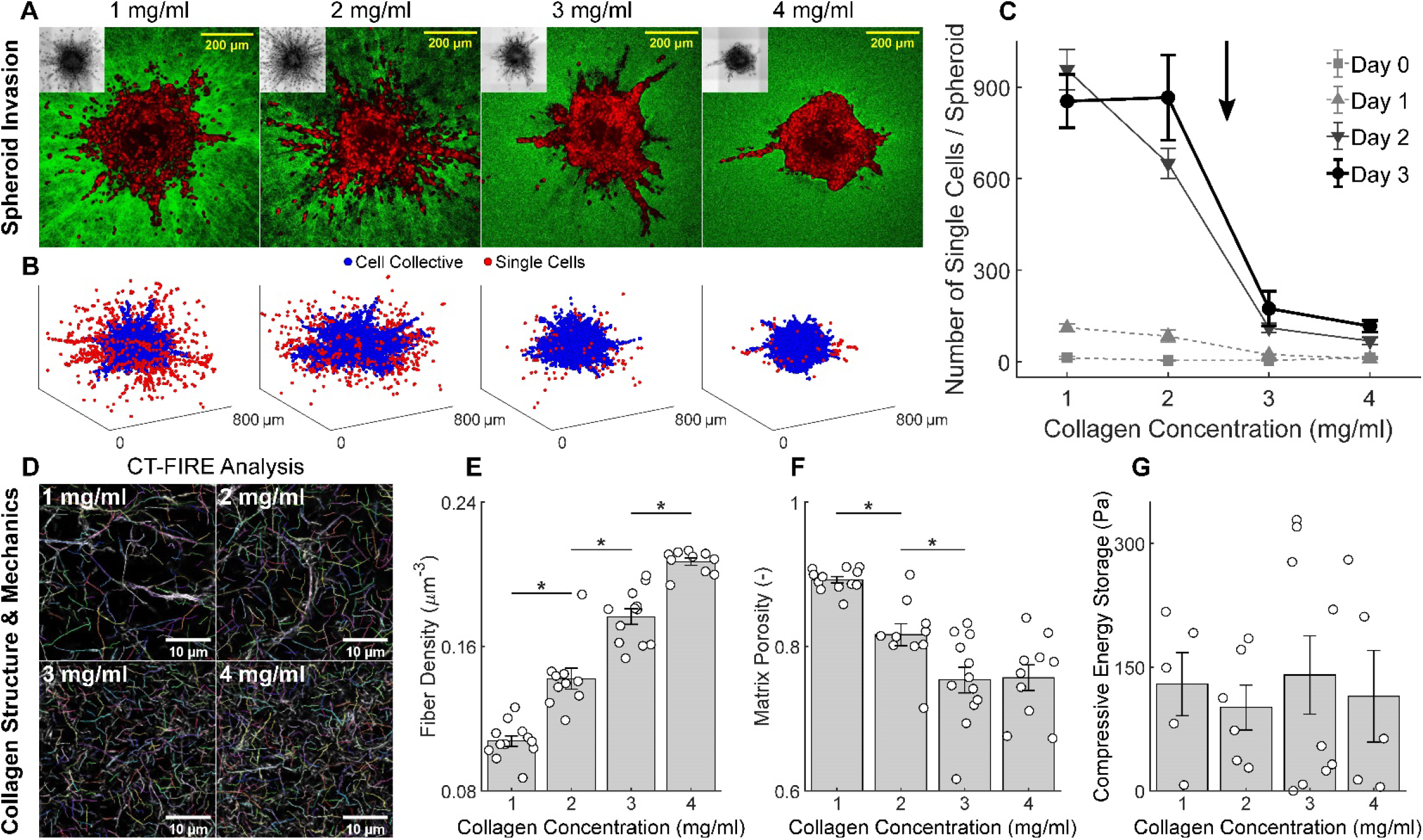
Collagen fiber density, not mechanics, modulates the switch in invasion phenotype observed in MDA-MB-231 spheroids. (**A**) Representative equatorial cross-sections of multiphoton images show MDA-MB-231 spheroids after 3 days of invasion in graded collagen concentrations (1 to 4 mg/ml) along with the associated DIC minimum intensity projections (insets). Single-cell migration is observed primarily in 1 and 2 mg/ml while collective migration is observed primarily in 3 and 4 mg/ml. (**B**) Corresponding 3D rendering of cell nuclei distributions identified from automated analysis of multiphoton image stacks (Methods). Nuclei are color-coded based on whether they remain within the cell collective (blue) or are detected as single cells (red). **(C)** Immediately after embedding in collagen (day 0), all cells are part of the multicellular collective with no invasion at any collagen density. As the spheroid evolves over time (days 1, 2 and 3), a striking gas-like phase and corresponding single cell escape progressively emerged at lower collagen concentrations (1 and 2 mg/ml) but not higher collagen concentrations (3 and 4 mg/ml). By day 3, a switch-like biphasic reduction in the number of single invading cells emerged when collagen concentration was increased from 2 to 3 mg/ml. The temporal evolution of single cell invasion as a function of collagen concentration supports the existence of criticality between 2 and 3 mg/ml, at which point the invasive phenotype switches abruptly from single to collective invasion. Single cell counting data are shown from days 0-1-2-3 and collagen concentrations of 1-2-3-4 mg/ml, n = 3 per group, except for day 0 - 1 mg/ml (n = 2) and day 2 - 2 mg/ml (n = 9). The significance of differences due to collagen concentration and time were quantified using a one-way ANOVA and post-hoc pairwise comparisons with Bonferroni correction. Statistical significance was achieved between 1 and 2 mg/ml at day 2 (p < 0.05), and between 2 and 3 mg/ml at days 1 (p < 0.05), 2 (p < 0.01), and 3 (p < 0.01), while no significant differences were observed between 3 and 4 mg/ml. The sharp and statistically significant transition between 2 and 3 mg/ml is therefore indicated with an arrow. We examined whether this transition is due to differences in collagen structure or mechanics. (**D**) High-resolution multiphoton images show representative acellular collagen networks at 1 to 4mg/ml, with individually segmented fibers from CT-FIRE analysis (*70*) as indicated by different colors. We quantified microstructural and mechanical properties of these collagen networks by combining multiphoton imaging and mechanical testing under confined compression (Methods and Supplementary Material). (**E**) Fiber density and (**F**) matrix porosity display clear trends with increasing collagen concentration between 1 and 4 mg/ml. (**G**) In contrast, compressive energy storage (under 18% compression), which quantifies nonlinear mechanical properties of the matrix, are indistinguishable between 1 and 4mg/ml. Therefore, collagen fiber density, rather than mechanics, represents a control variable for 3D cell jamming. Microstructural data are shown from 1 mg/ml (n = 12), 2 mg/ml (n = 10), 3 mg/ml (n = 12), and 4 mg/ml (n = 12) collagen gels. Mechanical data are shown from 1 mg/ml (n = 5), 2 mg/ml (n = 6), 3 mg/ml (n = 9), and 4 mg/ml (n = 5) collagen gels. * indicates statistical significance at p < 0.05.

Overall, these findings support the possibility of the existence of a critical collagen density at which MDA-MB-231 cells at the spheroid periphery transition in an almost switch-like fashion between distinct modes of invasion. Under lower matrix confinement, unjammed cells tend to invade individually as single cells or discrete cell clusters in a gas-like fashion. Under higher matrix confinement, however, these highly motile cells progressively slow at the spheroid periphery and within invading protrusions they tend to re-jam. For the invading cellular collective, these observations support the interpretation that high density collagen promotes steric hindrance and an associated confinement-induced jamming (*9, 39*). These events likely depend upon active remodeling of ECM by metalloproteases (*9*), cell generated traction forces (*40–42*), and the manner in which these tractions act to align collagen fibers (*41*). The invasive behaviors and material states observed from micro- and macro-spheroids of MCF-10A and MDA-MB-231 cells are summarized in Table 1.

**Table 1:** Across different spheroid systems, cell types, and experimental conditions, drastic changes in invasive phenotypes are reflected by transitions between material phases. Measured trends and observations are summarized relative to the early stage MCF-10A micro-spheroid. The two spheroid systems were labelled as “Micro” and “Macro” based on the approximate number of cells contained in the multicellular clusters. “Early Stage” and “Late Stage” conditions correspond to the early (day 3-5) and late (day 7-10) stages of evolution for MCF-10A micro-spheroid. “Low ECM Density” and “High ECM Density” correspond, respectively, to 2 and 4 mg/ml collagen concentration in which macro-spheroids were embedded. Regions within spheroids were roughly separated into core (“c”) and periphery (“p”) to compare trends in the measured values (Shape Index, Motility, Cell Volume). Such trends and the associated radial gradients underlie the observation of different material phases and invasive modalities. **\*** We note that within MCF-10A macro-spheroids in 4 mg/ml collagen, there was a radial increase in cell volume but not SI and motility. The column “Reference” specifies the figure where these observations are reported and discussed in detail in the following sections: ^**a**^*In the MCF-10A micro-spheroid, the core approaches a jammed, solid-like phase;* ^**b**^*In the MCF-10A macro-spheroid, the periphery invades as a locally unjammed fluid-like phase*; ^**c**^*In MDA-MB-231 macro-spheroids, the invasive phenotype switches abruptly as a function of ECM density.*

| Experimental Conditions |  |  | Measured Trends |  |  |  |  | Observations |  |  | Reference |
| --- | --- | --- | --- | --- | --- | --- | --- | --- | --- | --- | --- |
| *Spheroid System* | *Cell Type* | *Condition* | *Region* | *Shape Index* | *Motility* | *Cell Volume* | *Radial Gradient* | *Invasion* | *Invasive Modality* | *Material Phase* |  |
| **Micro**<br>$n_{cells} \sim O(10^1\text{-}10^2)$ | MCF-10A | Early Stage | c | - | - | - | No | No | - | Fluid | Fig. 1<sup>a</sup> |
|  |  |  | p | - | - | - |  |  |  |  |  |
|  |  | Late Stage | c | ↓ | ↓ | ↓ | Yes | Yes | Collective | Solid → Fluid |  |
|  |  |  | p | ↑ | ↑ | ↑ |  |  |  |  |  |
| **Macro**<br>$n_{cells} \sim O(10^3\text{-}10^4)$ | MCF-10A | Low ECM Density | c | - | - | - | Yes | Yes | Collective | Solid → Fluid | Fig. 2<sup>b</sup> |
|  |  |  | p | ↑ | ↑ | ↑ |  |  |  |  |  |
|  |  | High ECM Density | c | - | - | - | No* | No | - | Solid |  |
|  |  |  | p | - | - | ↑ |  |  |  |  |  |
|  | MDA-MB-231 | Low ECM Density | c | ↑↑ | ↑↑ | ↑ | No | Yes | Single Cell | Fluid → Gas | Figs. 3,4<sup>c</sup> |
|  |  |  | p | ↑↑ | ↑↑ | ↑ |  |  |  |  |  |
|  |  | High ECM Density | c | ↑↑ | ↑↑ | ↑ | Yes | Yes | Collective | Fluid → Solid |  |
|  |  |  | p | ↓ | ↓ | ↓ |  |  |  |  |  |

### Mapping a hypothetical jamming phase diagram

Mechanical interaction between the invading cellular collective and the surrounding ECM is impacted by a variety of factors, including but not limited to fiber composition, connectivity, stiffness, porosity, nematic alignment, cell-matrix adhesion, matrix proteolysis, cellular and nuclear stiffness, contraction, and matrix deformation (*1, 39, 42–46*). Among the most primitive of these interactions is mutual volume exclusion, where cells, or cells and ECM fibers, cannot occupy the same space at the same time. This physical limitation is related to geometric confinement and steric hindrance, wherein motion of self-propelled cells can be constrained by geometry (*9, 39, 47*). Rather than exhaustively capturing all the aforementioned interactions, we took a minimalist approach to address a specific question. Based on our experimental findings, we asked if we can recapitulate the observed migratory phenotypes by varying only two key biophysical factors: cellular propulsion and ECM density. To answer this question, we developed the minimal 2D model that characterizes in-plane cell and ECM interactions, and captures the behavior of a dense cellular collective comprising the early tumor and its invasion into a dense, but porous, ECM.

In the model, cortical tension and cell elasticity were incorporated much as in traditional vertex models (*14, 25*). However, we modified those previous models through combination with an agent-based approach (*48*) that takes into account physical interaction between cells and the ECM, as well as the possibility of single cell detachments. Each cell was assigned an elastic response to departures from a preferred area and a viscoelastic response to departures from a preferred perimeter (Supplementary Methods). In addition, each cell was endowed with self-propulsion of magnitude v_0,_ which acts as a vector with randomly generated polarity (*26*). Adjacent cells were given their own cell boundaries that moved along with the cell to which it belongs. To highlight the roles of steric hindrance and system geometry, ECM density was modeled by tuning the spatial density of matrix fibers, which are represented as randomly distributed discrete posts that are fixed in space and do not adhere to cells. Despite these simplifications, our 2D hybrid model shows a remarkably rich repertoire of dynamical behaviors and captures well the striking phenomena reported in the experimental models (Fig. 5).

**Fig. 5.**
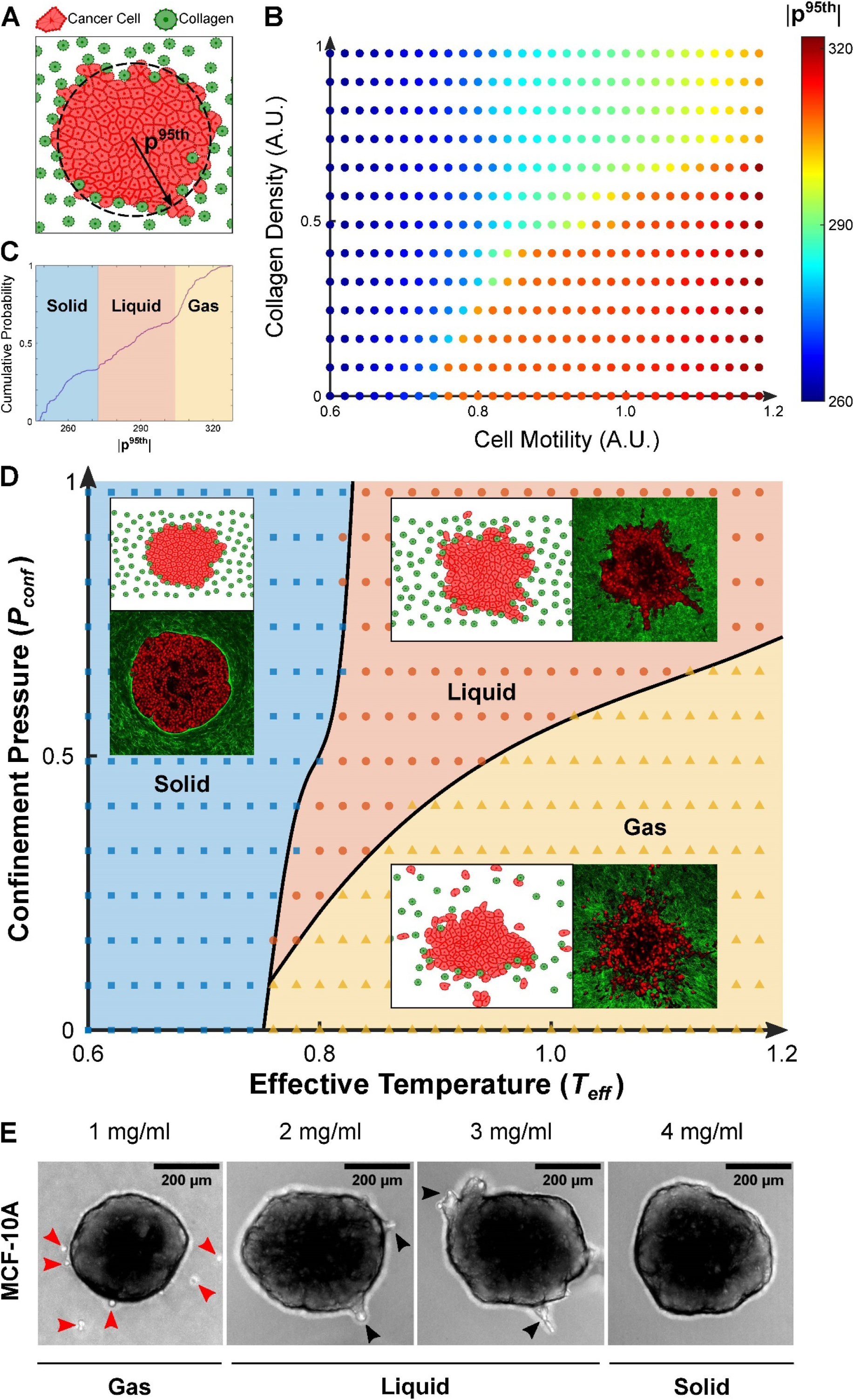
A 2D computational model of a multicellular cluster in collagen reveals that tumor invasion phenotypes and associated material states are governed by a jamming phase diagram. **(A)** The hybrid computational model of tumor invasion into ECM is characterized by cancer cells (orange particles) that can move in random directions with varying degrees of self-propulsion (Supplementary Methods). At the beginning of each simulation, cancer cells are organized to form a circular collective that is surrounded by collagen (green particles), arranged randomly and with varying spatial densities (Supplementary Methods). At the end of each simulation, the 95^th^ percentile of the radial cell positions (**p**^95th^) with respect to the centroid of the collective is used as a read-out of the degree of invasion. **(B)** A diagram is generated by gradually incrementing two state variables: cell motility and collagen density, both expressed in arbitrary units (A.U.). Data points are color-coded according to the mean value of **p**^95th^ over n = 10 simulations, each corresponding to randomly assigned positions of the collagen particles and orientations of the cell motility vectors. Three notable regions can be distinguished in the diagram and qualitatively correspond to solid-, liquid-, and gas-like behaviors at the invasive front (Movies S2-4). **(C)** These three regions can be distinguished from distinct elbow regions in the cumulative probability distribution of **p**^95th^ generated from all simulations. We identified the 34^th^ and 64^th^ percentiles as robust thresholds (cf. Table S2) to separate solid from liquid and liquid from gas phases, respectively. **(D)** The resultant map represents a jamming phase diagram, now color-coded to indicate solid-like (blue squares), fluid-like (orange circles), and gas-like (yellow triangles) material phases. In analogy with equilibrium thermodynamic systems, here cell motility is replaced with an effective temperature (*T_eff_*, Box 1) while collagen density is replaced with a confinement pressure (*P_conf_*, Box 2). By tuning only two state variables, the model recapitulates much of the experimentally observed behaviors. For each material phase on the diagram, representative multiphoton images from experiments are shown in comparison to representative computational snapshots (insets). In the solid-like phase (blue area), a non-invasive MCF-10A spheroid in high collagen density (4 mg/ml) is shown in comparison to the result of a simulation parameterized with low cell motility (0.2) and high collagen density (0.82). In the fluid-like phase (orange area), an MDA-MB-231 spheroid collectively invading in high collagen density (4 mg/ml) is shown in comparison to the result of a simulation parameterized with high cell motility (1.0) and high collagen density (0.82). Finally, in the gas-like phase (yellow area), an MDA-MB-231 spheroid scattering into single cells in low collagen density (2 mg/ml) is shown in comparison to the result of a simulation parameterized with high cell motility (1.0) and low collagen density (0.21). Overall, we observe that at low cell motility, and thus low *T_eff_*, the system is homogeneously “cold” and the spheroid shows a non-invasive, solid-like behavior regardless of collagen density. However, at higher *T_eff_* the collagen density, and hence *P_conf_*, determines fluid-like or gas-like behaviors. Phase boundaries (black lines) on the jamming phase diagram are obtained as best-fit curves that separate data points belonging to different material phases. Unlike traditional thermodynamic phase transitions, where the boundary lines mark clear transitions between material phases, in our cellular systems material transitions are continuous and smeared. Thus, the boundary lines mark regions of coexistent phases, where near each phase boundary, the material phases become indistinguishable. The proposed diagram also predicts the existence of a “triple point” where solid-, liquid-, and gas-like phases coexist, and below which direct solid-to-gas transitions occur. **(E)** To test the plausibility of such prediction we ran an invasion assay in graded collagen concentrations (1 to 4 mg/ml) using MCF-10A spheroids. which, according to the phase diagram, are characterized by a lower *T_eff_* with respect to their MDA-MB-231 counterparts. The periphery of MCF-10A spheroids was found to remain solid-like and non-invasive in 4 mg/ml, to fluidize and invade collectively in 3 and 2 mg/ml and, more importantly, to separate directly into individual gas-like cells in 1 mg/ml. These findings support the direct individualization of cancer cells from a nearly jammed tumor as predicted by our jamming phase diagram.

We assessed the degree of invasion at the end of each simulation based on the 95^th^ percentile of the radial cell position (**p**^95th^) with respect to the cluster centroid (Fig. 5A). By modeling the behavior of cell clusters while varying both cell motility and collagen density, we found that the resulting map of **p**^95th^ (Fig. 5B) depicts three notable regions, each corresponding to a distinct invasion phenotype (Fig. 5C-D). Specifically, low values of **p**^95th^ correspond to a low degree of invasion (blue area in Fig. 5C-D, Movie S2), intermediate values of **p**^95th^ are associated with collective cell invasion (red area in Fig. 5C-D, Movie S3), while high values of **p**^95th^ are associated with single cell invasion (yellow area in Fig. 5C-D, Movie S4). Using the cumulative probability distribution of **p**^95th^, we mapped these three regions onto a jamming phase diagram (Fig. 5C-D, Table S2). Within this phase diagram solid-like, fluid-like, and gas-like phases dominate the invasion of the simulated spheroid in a manner that is remarkably similar to the invasive phenotypes observed experimentally (Fig. 5D).

This computational model not only recapitulated our experimental observations, but further suggested that a cancer cell collective can invade into surrounding ECM by following a variety of paths on the proposed jamming phase diagram. When cell motility (v_0_) is small the cellular collective shows solid-like behavior with little migration or invasion, and thus resembles nearly jammed MCF-10A spheroids in high density collagen (Fig. 5D). When v_0_ is progressively increased while keeping collagen density fixed and high, a fluid-like cellular collective flows in branches that protrude from the continuous tumor mass much alike the MDA-MB-231 spheroids undergoing a jamming transition in high collagen density (Fig. 5D). When v_0_ is held fixed and collagen density is decreased, single cells and cell clusters detach from the collective thus mimicking the gas-like dispersion observed in MDA-MB-231 spheroids undergoing fluid-to-gas unjamming in low collagen density (Fig. 5D). Furthermore, when both v_0_ and collagen density are low the phase diagram predicts a “triple point”, where solid-like, fluid-like and gas-like phases coexist. This physical picture implies a novel invasion modality, namely, a solid-to-gas transition in which cancer cells detach from the nearly jammed spheroid and individualize. We tested the plausibility of such prediction from the phase diagram by performing an invasion assay using MCF-10A spheroids in graded collagen concentrations (1 to 4 mg/ml). Over 48 hours in 4 mg/ml collagen, MCF-10A spheroids remained solid-like. As extracellular confinement progressively reduced until collagen density reached 1 mg/ml, we observed direct detachment of single MCF-10A cells from the main spheroid, and migration as gas-like particles (Fig. 5E and Movie S5). We suggest that the numerous variables that govern collective behavior may combine so as to reduce to only two overriding variables, namely, an effective temperature (Box 1) and an effective confinement pressure (Box 2). Overall, we conclude that the proposed jamming phase diagram provides a useful guide for thought and, potentially, a unifying mechanistic interpretation of jamming/unjamming transitions in cancer invasion.

## DISCUSSION

The principle finding of this report is that transitions in tumor cell morphology, packing, migration, and invasion are governed by coexisting material phases. These include a jammed solid-like phase, and unjammed fluid-like and gas-like phases. We show that, depending on cell and matrix properties, the tumor mass can invade collectively either by undergoing unjamming, as in the case of MCF-10A spheroids in 2 mg/ml collagen, or by means of jamming, as in the case of MDA-MB-231 spheroids in 4 mg/ml collagen. This behavior suggests that tumor invasion is not the sole result of progressive tissue fluidization, but rather is a richer process wherein a variety of routes towards phase separation allow the tumor to attain different invasive phenotypes. Within the experimental systems employed here, our findings identify cell motility and collagen density as key state variables controlling the phase separation of tumor cells *en masse,* governed by an overriding jamming phase diagram that was derived computationally (Fig. 5D). Although this phase diagram expresses the same material phases of the diagram hypothesized by Ilina et al. (*15*), it differs from theirs in several aspects, and is therefore novel. For example, it differs from Ilina et al. in the types of phase transitions that a tumor can undergo. In addition to the liquid-to-solid, solid-to-liquid, and liquid-to-gas transitions observed in invading tumor spheroids (Figs. 1-4), this novel phase diagram also predicts a distinctive solid-to-gas transition – reminiscent of sublimation – which we confirmed experimentally (Fig. 5E). In addition, we suggest that the many physical factors that characterize tumor cells, their interactions with one another, and their interactions with the ECM, may be incorporated to extend the proposed phase diagram and map into a small set of ‘effective’ thermodynamic variables, such as an effective temperature (Box 1) and an effective confinement pressure (Box 2).

As used here, the phrase ‘coexistence’ of material phases has two distinct but interrelated connotations.As suggested previously (*49*), in the vicinity of a phase boundary, seemingly modest changes of cellular or ECM properties may have the potential to precipitate striking changes of material phase and invasion phenotype. The second connotation suggests that in the same spheroid the cellular collective can express macroscopic regional differences, such as a solid-like core coexisting with a fluid-like invasive branch. In MCF-10A spheroids, for example, cells at the periphery compared with cells near the core tend to be systematically more elongated, more variable in shape, and more migratory (Figs. 1 and 2). In the context of active force fluctuations and associated metabolism, cells at the periphery are also more dynamic (*16, 50*). Conversely, for post-metastatic MDA-MB-231 spheroids, cells at the core are larger, more variable in shape, and more migratory with respect to MCF-10A counterparts, while the periphery is highly sensitive to changes in collagen density (Figs. 3 and 4). Compared to MCF-10A cells, they also generate higher traction forces (*51*) and are more metabolically responsive (*52*). Together, this constellation of structural, migratory, mechanical, and metabolic factors is consistent with the existence of an effective temperature, *T_eff_*, that is spatially heterogeneous. Such a physical picture would help to explain, and perhaps to generalize, the x-axis of the hypothesized jamming coexistence phase diagram (Fig. 5D; Box 1). As regards the y-axis of the hypothesized phase diagram, it is well established that both solid and fluid stresses within the spheroid core are compressive, and that cellular and nuclear volumes in the core are reduced, as if under compression (*16, 22*). Such a compressive state of stress is thought to arise in part from cellular proliferation under the constrain to ECM confinement (*53*). In addition, osmotic pressure decreases cellular volume, increases cell stiffness and thereby decreases invasiveness of peripheral cells (*16*). It remains unclear, however, how the solid stress due to both neighboring cells and ECM, interstitial fluid stress, and osmotic stress combine to generate the hypothesized confinement pressure *P_conf_*. Such connections, if they could be established, would help to explain, and perhaps to generalize, the y-axis of the hypothesized jamming coexistence phase diagram (Fig. 5D; Box 2).

A central role of the EMT in tumor cell motility, invasiveness, and metastasis, is well established but has recently become a point of contention (*54*). Our results suggest that relative to EMT, jamming and unjamming transitions represent a complementary route for establishment of cell motility and invasiveness. Indeed, in at least one system unjamming and collective migration are known to occur in the absence of EMT (*18*). In the 3D systems examined here, MDA-MB-231 cells lack E-cadherin and are therefore mesenchymal-like (Fig. S1). However, when embedded in high density collagen, these cells undergo confinement-induced jamming while still retaining fluid-like collective invasive behavior (Fig. 3) (*9*), thus indicating that jamming and unjamming transitions are highly influential in collective behavior. Establishing the extent to which cell jamming/unjamming may be complementary to or, potentially, may subsume EMT in directing collective cellular motion seems a necessary step for fuller understanding cancer progression.

Compared to inert materials, cellular collectives are biologically active and displaced far from thermodynamic equilibrium. Thus, the collective material phases identified here, and the associated jamming/unjamming transitions, are not to be confused with first order or second order transitions occurring in systems close to thermodynamic equilibrium. Like the glass transition (*55*), jamming/unjamming transitions generally display a discontinuity in the number of contacts, which are characteristic of first-order phase transitions, but display diverging correlation length scales, which are characteristic of second-order phase transitions (*14, 56, 57*). Similarly, we find that cell migration shows smooth radial changes in invading tumor spheroids (Figs. 1 to 3) but sharp transitions as a function of collagen concentration (Fig. 4). As opposed to a phase transition that is binary and sharp, as might occur in an equilibrium system, the transition between a jammed and an unjammed cellular phase is continuous and smeared both in space and in time (*13, 16*). Just as there is no latent heat and no structural signature of melting for inert materials approaching the glass melting point, so too in invading tumor spheroids there is no sharp transition in cellular shapes or migration speeds. Therefore, the material phase of a cellular collective needs to be defined via functional terms –cellular migratory persistence, cooperativity, a target shape index, and cellular migratory propulsion (*14, 25, 58*).Yet, despite these differences, the cell collectives herein are observed to transit various material states within a phase diagram that bear superficial resemblance to that of common thermodynamic systems. These results suggest that collective cellular migration, invasion and escape from a cellular mass involve biophysical processes far richer than previously anticipated, but may be governed by basic physical principles. We have shown how specific cell and ECM properties can be reduced to a set of ‘effective’ thermodynamic variables describing the material phase of the invasive cell collective, and thus mapping a jamming phase diagram of tumor invasion. How the local material phase of the cellular collective and its mechanical properties might impact the emergence of driver mutations remains unknown. Deformation of the cell and its nucleus associated with migration within a highly confining microenvironment is known to cause loss of nuclear envelope integrity, herniation of chromatin across the nuclear envelope and DNA damage (*59*), but the impact of cell and nuclear elongation in connection with unjamming remains unstudied. Conversely, how driver mutations and resulting subclonal heterogeneities might impact the local material phase is also unclear. When such interactions become appreciable, tumor dynamics would then be seen to be a multifaceted problem in mechanogenetics (*60*).

## MATERIALS AND METHODS

### Cell lines and culture media

Non-tumorigenic MCF-10A and metastatic MDA-MB-231 breast epithelial cell lines were purchased from American Type Cell Culture Collection (ATCC) and cultured using standardized media and conditions (*16, 40*). MCF-10A cells were cultured in DMEM/F-12 (ThermoFisher, No. 11330032) supplemented with 5% horse serum (Invitrogen, No. 16050122), 20 ng/ml EGF (Peprotech, AF10015; ThermoFisher, No. 10605HNAE), 0.5 mg/ml hydrocortisone (Sigma-Aldrich, No. H0888), 100 ng/ml cholera toxin (Sigma-Aldrich, C8052), 10 μg/ml insulin (Sigma-Aldrich, No. I1882). MDA-MB-231 cells were culture in DMEM (Corning, No. 10013CV), supplemented with 10% fetal bovine serum (ATCC, No. 302020). Both media recipes contained 1% penicillin/streptomycin (ATCC, No. 302300; ThermoFisher, No. 15140122). Cells were maintained at 37°C and 5% CO_2_ in a cell culture incubator.

### Spheroid formation

Two distinct protocols were used to generate the micro- and macro-spheroid models used in this study. First, micro-spheroids were formed by trypsinizing and embedding MCF-10A cells within an interpenetrating network (IPN) consisting of 5 mg/ml Alginate (FMC Biopolymer) and 4 mg/ml Matrigel (Corning, No. 354234) as previously shown (*16, 19*). Cells were mixed with the gel precursor solution, which was allowed to gel inside an incubator before adding culture media. The shear modulus of the double network can be tuned via calcium cross-linking and was herein set to 300 Pa to reproduce the stiffness of malignant breast tissue (*19*). Within the IPN, cells proliferated to form micro-spheroids that began invading into the gel after approximately 7 to 10 days in culture. Second, MCF-10A and MDA-MB-231 macro-spheroids were generated by seeding approximately 10^3^ cells in each of the 96 wells of an ultra-low attachment plate (Corning, No. 07201680) and allowed to form for 48 hours in presence of 2.5% v/v Matrigel. We verified that addition of a small volume fraction of Matrigel allows formation of MDA-MB-231 spheroids, which would otherwise form only loose aggregates (Fig. S6) (*38*). MCF-10A spheroids were formed under the same conditions to ensure consistency. Once formed, individual spheroids surrounded by a small volume of media were transferred in microwells (10 mm in diameter) inside of glass bottom 6-well plates (MatTek, No. P06G-0-10-F) by pipetting 5 μl drops on each of the coverslips. Each spheroid was covered by 145 μl of ice-cold, rat-tail collagen I solution to achieve a total volume of 150 μl and a specific collagen concentration in each microwell. Collagen solutions were prepared by mixing acid-solubilized collagen I (Corning, No. 354249) with equal volumes of a neutralizing solution (100 mM HEPES buffer in 2x PBS) (*61*). The desired collagen concentration was reached by adding adequate volumes of 1x PBS. Collagen solutions at different concentrations (1, 2, 3, and 4 mg/ml) polymerized for 1 hour at 37°C. The cell culture plates were rotated every minute for the first 10 minutes of polymerization to guarantee full embedding of the spheroid within the 3D collagen matrix. Finally, 2 ml of culture media was added and the 3D organotypic culture was placed inside the incubator for a variable amount of time. Media was refreshed every two days.

### Measurement of micro-spheroid dynamics

MCF-10A cells were stably transfected with GFP-linked nuclear localization (NLS-GFP). To track the movement of individual cells, micro-spheroids at early (3-5 days) and late (7-10 days) stage of growth were imaged every 10 minutes for 12 hours in a customized incubator (37°C, 5% CO_2_, and 95% humidity) on a confocal microscope (Leica, TCL SP8), with acquisition of fluorescent and bright-field image z-stacks (Fig. 1A-B). Analysis of cell migratory dynamics was done based on the obtained image z-stacks according to the procedures outlined below, implemented in custom MATLAB programs (cf. *nuclei identification*). Using an adaptation of previously published methods (*62*), the 3D coordinates of each cell nucleus were identified at each frame based on maximum intensities from the fluorescent images. Cell trajectories were constructed through minimization of overall nuclear displacements between sequential frames. Cell positions from each time point were aligned to the micro-spheroid geometric center at the initial time point *t*_0_ to account for any translational motion that may have occurred due to stage drift. To separate the contribution of Coherent Angular Motion (CAM) of the early micro-spheroid from migratory dynamics due to individual cell rearrangement, we solved for the rigid rotational transformation *ω* that best reproduced the changes occurred in cell nuclei positions (*p*(*x*, *t*)) between z-stacks during the time interval Δ*t*. *ω* is defined as the rotation that minimized the metric *d*(*ω*|*t*) = ‖*p*(*x*, *t* + Δ*t*) − *R*(*ω*Δ*t*)*p*(*x*, *t*)‖. This rigid rotational motion from CAM is removed from cell trajectories before further calculations of cellular dynamics. Root mean squared (RMS) speeds for each cell were calculated using the instantaneous velocity vectors. Mean squared displacements (MSD) were computed as a function of time interval, *MSD*(Δ*t*) = 〈|*p*(*x*, *t* + Δt) − *p*(*x*, *t*)|^2^〉. Here, 〈…〉 denotes an average over time, with overlap in time intervals.

### Measurement of macro-spheroid dynamics

A customized spinning disk confocal setup equipped with an environmental chamber (37°C, 5% CO_2_, 80% relative humidity) was used for imaging spheroid invasion in collagen. For each spheroid within a 6-well plate, differential interference contrast (DIC) images were collected every 10 minutes for 48 hours as 3×3 tiled, 200 μm z-stacks. A 10x air objective was used to image large areas (1450.3 × 1450.3 μm) with a resolution of 1.126 μm/pixel. Automated stitching based on global optimization of the 3D stacks (*63*) was carried out using ImageJ (v. 1.52g, National Institute of Health, Bethesda, MD). Minimum intensity projections of the stitched DIC time-lapse data were used to visualize the 3D data sets as 2D movies while maximizing the image contrast (dark spheroid/cells on a light background). These DIC movies were imported in MATLAB (R2019a, Mathworks, Natick, MA), where images from individual time frames were down-sampled by 50% (2.252 μm/pixel) to optimize the processing speed. De-jittering of movies was achieved using an optimized image registration algorithm (*64*). Temporal evolution of the projected macro-spheroid area was segmented using Otsu’s method, where the optimal threshold was calculated over three consecutive frames to smooth changes in area between frames. The resulting binary mask allowed us to separate between the spheroid interior (i.e., main spheroid) and exterior (i.e., single cells migrating in collagen, if any). To estimate the direction and speed of collective migration within the macro-spheroid, we employed Farneback’s optical flow method using the Matlab function *estimateFlow* with the option *opticalFlowFarneback* and a Gaussian filter size of 8 pixels while keeping all the other parameters to their default values. At each frame, velocity vectors are obtained for seed points spaced 2 μm apart that are inside the spheroid mask. To account for the effect of collective motion from spheroid growth on migratory dynamics, we estimated spheroid radial growth rate through tracking of change in spheroid mask area over time. This velocity was interpolated for each seed depending on its location within the spheroid, and were removed from the velocity vectors before RMS speed calculations Single cells lying outside the spheroid mask were detected based on local intensity minima at each frame, and tracks constructed by minimizing displacements of cells between consecutive frames. Both collective and single cell velocity vectors were smoothed with a moving average filter of 3 frames with unity weighting in the temporal domain. RMS speed were calculated for each cell/seed using the instantaneous velocity vectors.

### Multiphoton microscopy

In order to image cell nuclei within large spheroids, we adapted a technique known as CUBIC (Clear, Unobstructed Brain/Body Imaging Cocktails) which was originally developed to enable optical clearing and high-resolution imaging of murine organs (*28*). Briefly, CUBIC employs hydrophilic reagents to remove lipids (the main source of scattering within tissues), while preserving fluorescent proteins. Following the original protocol by Susaki et al.(*28*), we prepared a mixture of 25% wt urea (Fisher Scientific, No. U15), 25% wt Quadrol (N,N,N′,N′-Tetrakis(2-hydroxypropyl)ethylenediamine, Sigma-Aldrich, No. 122262), 15% Triton X-100 (Sigma-Aldrich, No. T8787), and dH_2_O. At regular intervals after embedding, spheroids were fixed overnight using cold 4% PFA (Fisher Scientific, No. AAJ19943K2), and washed out twice using 1x PBS. Samples were pretreated for 2 hours in 2 ml of ½ diluted CUBIC reagent (50% vol/vol dH_2_O) and then immersed in 1 ml CUBIC reagent with 2 μM DAPI (Fisher Scientific, No. D1306) under gentle shaking at room temperature. The CUBIC reagent and DAPI were refreshed every two days and samples were cleared for up to 14 days prior to imaging. The 3D organization of optically cleared spheroids in collagen was imaged using a Bruker Ultima Investigator multiphoton microscope (MPM). The laser beam was focused onto the spheroids through a 16x water-immersion objective (Nikon, 0.8 N.A., 3 mm working distance) mounted in upright configuration. We used an excitation wavelength of 880 nm to image both invading spheroids and collagen. Two-photon excitation fluorescence (TPEF) from DAPI stained nuclei was collected through a bandpass filter centered at 550 nm with a bandwidth of 100 nm, while second harmonic generation (SHG) signal from the collagen matrix was collected through a bandpass filter centered at 440 nm with a bandwidth of 80 nm. Images were collected using 1024 × 1024 pixels at a resolution of 0.805 μm/pixel and a pixel dwell time of 10 μs. Stacks were acquired using 5 μm steps and a thickness (variable in the range 400-1000 μm) that was determined depending on the spheroid size and degree of invasion. Laser power and photomultiplier tube voltage were increased to maintain a nearly constant signal across the spheroid and interpolated linearly though the PraireView software during acquisition.

### Cell shape characterization

We used a customized analysis procedure in MATLAB to characterize the cell shapes from both micro- and macro-spheroids. Steps in the analysis pipeline are illustrated in Fig. S2 and described below.

#### Nuclei identification

Nuclei centers were identified based on local intensity maxima from fluorescent image stacks (DAPI or GLP-NLS) with adaptations of previously published methods (*62, 65*). In brief, images were preprocessed with MATLAB function *medfilt2* and *smoothn* to minimize background noise and smooth out intensity variation within each nucleus. To reduce noise, background intensity outside the spheroid were removed by thresholding for the lowest 10% of signal intensities. Local intensity maxima were then identified from each 2D image slice, and a connected component analysis and a search radius of 5 μm in 3D was used to identify nuclei centers. Accuracy of nuclei identification of each spheroid was assessed via visual inspection of the overlay between identified nuclei positions with fluorescent nuclei image stacks (Fig. S2B). Further validation of this nuclei identification algorithm was carried out by comparing algorithm-identified cell counts with the manual counts obtained using a hemocytometer after spheroid dissociation via trypsinization. Validation was performed for macro-spheroids seeded at different sizes for both MCF-10A and MDA-MB-231 cells (Fig. S2G).

#### Spheroid boundary segmentation

Spheroid boundaries were identified from either bright-field microscopy (for micro-spheroids) or multiphoton SHG (for macro-spheroids) z-stacks. The spheroid boundary was identified in a two-step (coarse and fine) segmentation process, adapting previously published protocols (*66, 67*). Image z-stacks were preprocessed with intermediate steps including *histogram equalization*, *wiener* (8×8 pixels) and *median* (5×5 pixels) filters to enhance image contrast. Initial coarse segmentation was performed using the watershed algorithm on the gradient image of the resulting image z-stacks, supplemented with nuclei position information. The fine segmentation step involved adaptive k-means clustering based on variation in pixel intensities in the output image from the coarse segmentation step. The number of clusters segmented was determined automatically based on the distance between new and existing cluster centers from the previous iteration. The final spheroid boundary was taken as the data cloud outline of the spheroid cluster that formed a connected region with the largest volume. Validation was done by visual inspection of the overlay between identified spheroid boundaries with the raw image z-stacks (Fig. S2C).

#### Bounded Voronoi Tessellation

Voronoi tessellation of the nuclei centers were based on the observation by Voigt and Weis (*68*), who have shown that the vertices of each Voronoi cell are solution to sets of linear inequalities indexed by their nuclei centers. Delaunay triangulation of the nuclei centers was used to identify the Voronoi neighbors for each cell. The spheroid boundary data cloud was then downsampled to 5% while maintaining the shape of the spheroid. The set of linear inequality for each cell was then constructed from the union between the perpendicular bisectors between edges connecting the cell and its Voronoi neighbor; and the spheroid boundary identified from segmentation. Some cell-free regions in the macro-spheroid core caused the Voronoi tessellation to give falsely large cell volumes that included these regions, a double thresholding was applied to discard these cells. Voronoi cells were discarded if cell volumes exceeded 3000 μm^3^ or maximum distance of a cell to its immediate neighbors was in the top 5% of average maximum distance between all neighbors (Fig. S2F).

### Immunofluorescence

MCF-10A and MDA-MB-231 cells were cultured as a 2D monolayer on glass bottom plates for 6 days. Cells were fixed using either 4% PFA (20°C for 30 min) or Methanol (−20°C for 30 min), permeabilized with PBS / 0.2% Triton X-100 (20°C for 10 min), followed by incubation with primary antibody diluted in PBS / 10% goat serum / 1% BSA / 0.2% Triton X-100 (4°C overnight), rinsed, and incubated with secondary antibody diluted in PBS / 10% goat serum / 1% BSA / 0.2% Triton X-100 (20°C for 1 hour), and counterstained with 2nM DAPI (20°C for 10 min). Cells were rinsed and stored in 1x PBS for confocal fluorescence microscopy. Imaging was performed using an Olympus FV3000 confocal microscope equipped with a 20x/0.75N.A. objective. Image stacks from all samples are shown as maximum intensity projections. Proteins of interest were detected with the following primary antibodies from Cell Signaling Technologies: E-cadherin (1:200, #3195), Vimentin (1:100, #5741). Fluorescent labeling was carried out using the following secondary antibodies from Thermo Fisher Scientific: goat anti-rabbit AlexaFluor 488 (1:500, A-11008), goat anti-rabbit AlexaFluor 594 (1:500, A-11012).

### Western blotting

Lysates from MCF-10A and MDA-MB-231 monolayers were collected into ice-cold RIPA buffer with protease inhibitor cocktail and processed by incubation on ice for 30 minutes with sonication halfway through. Lysates were cleared by centrifugation at 13000g for 5 minutes and the supernatant was collected. Lysates were mixed with equal volume Laemmli 4x sample buffer (BioRAD) with 1M DTT and boiled for 6 minutes. Proteins of interest were detected using standard Western blotting with the following antibodies, all from Cell Signaling Technologies: E-cadherin (1:10,000, #3195), N-cadherin (1:1000, #13116), Vimentin (1:1000, #5741), GAPDH (1:10,000, #5174).

### Collagen gel structure and mechanics

Acellular collagen gels were polymerized within cylindrical PDMS wells. Structural and mechanical characterizations were carried out by using previously published methods (*69*). Briefly, high resolution multiphoton microscopy and analysis via CT-FIRE, and open-source code for segmentation of collagen fibers (*70*), led to measurement of matrix porosity and fiber density. Uniaxial confined compression experiments and fitting of experimental data to a finite deformation model for biphasic mixtures, led to estimation of bulk material properties and fluid transport properties (*71*). Due to the nonlinear mechanical behavior of collagen, we report the elastic energy storage under 18% compression as a measure of material properties and the hydraulic permeability from Darcy’s law as a measure of fluid transport properties. Further information can be found in the Supplementary Methods.

### Agent-based modeling

We developed a computational model that represents a hybrid between vertex(*25*) and particle-based(*48*) approaches. On a 2D computational domain, 152 cells were generated by a Voronoi tessellation of points randomly seeded (with Poisson sampling) in a circular area to form a cellular collective. Individual cells within the cellular collective were endowed with a randomly oriented cell motility vector v_0_, viscoelastic mechanical properties, and cell-cell interactions described in the form of a Lennard-Jones potential (Equation S7). Overall, the cellular collective was modelled as a confluent tissue, with no birth and death events, and was considered homogenous (in terms of cell properties such as adhesion and inherent motility). Collagen was represented as posts that are randomly distributed around the cell collective and was modeled as non-motile and rigid, and thus confining the cells. (Supplementary Methods). To capture all possible invasion patterns, the model was configured with gradual increase in the magnitude of the cell motility vector v_0_ and the density of collagen particles, which were chosen as state variables, while keeping constant all other simulation parameters (Supplementary Methods). To account for the random orientation of the motility vector and for the randomness in spatial configuration of collagen particles, we ran 10 simulations (n=10) for each pair of state variable configuration in the model, and used the averaged result to represent the cell collective behavior for that state variable combination. At the end of each simulation, the mean distance of the outermost 5% cells from the cellular collective centroid was used to represent the overall invasive behavior of the cellular collective. This measure was then pooled from multiple simulations across the range of values varied for the two state variables, and generated a histogram distribution used to identify the threshold criteria that best identified the invasive phenotypes and material phases: outermost distances below the 34th percentile were labeled as solid-like, between the 34th and 64th percentiles were labeled as fluid-like, and above the 64th percentile were labeled as gas-like. Within the model, these different material phases indicated different spheroid behaviors: a solid-like behavior represents lack of invasion, a fluid-like behavior represents collective invasion, and a gas-like behavior represents single cell invasion. Mapping the material phases and invasion patterns for each state variable combinations resulted in the jamming phase diagram. Sensitivity analysis showed the choice of exact threshold criteria, within the examined range, did not significantly impact configuration of the resultant phase diagram (Table S2). Additionally, mean cell shape (defined as 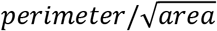 for cells in 2D), for cells in the solid-like phase were smaller (p<0.001) than cells in the fluid-like phase. Further information can be found in the Supplementary Methods.

### Statistical analysis

All of the data was analyzed in MATLAB. Experimental data are presented as mean ± STD. One-tailed t-tests were used to assess differences between spatial regions (core vs periphery) and stages of evolution (early vs late) of the micro-spheroid. A one-way ANOVA was used to test differences due to collagen concentration in the macro-spheroid, and post-hoc pair-wise comparisons were performed using the Bonferroni correction. Unless otherwise stated, p<0.05 was considered statistically significant.

#### Maximum likelihood estimation (MLE)

We fitted SI distributions to the *k-gamma distribution* using maximum likelihood estimation as described previously (*26*). In brief, for a data set {*x*_*i*_}*i* = 1,…, N, the likehood function to fit is 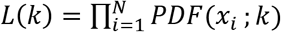, where PDF(*x*; *k*) = *k*;^*k*^*x*^*k*−1^*p*^−*kx*^/Γ(*k*) is the *k-gamma* probability density function. For our purposes, the SIs were shifted and normalized, and defined as 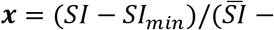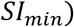, with *SI*_*min*_ = 5.413 based on Merkel and Manning (*24*).

###### Box 1: Effective Temperature

The standard definition of temperature is well understood, but it is sometimes useful to define an ‘effective temperature’ that has a functional equivalence. For example, Edwards and Oakeshott (*72*) in a conceptual leap considered the physics governing a powder, which is representative of a wider class of an inanimate inert collective systems that includes sand piles, pastes, colloid suspensions, emulsions, foams, or slurries (*73*). In such a collective system, thermal fluctuations are insufficient to drive microstructural rearrangements. As a result the system tends to become trapped far from thermodynamic equilibrium (*73*). Edwards and Oakeshott suggested that this class of collective systems might be understood in terms of the statistical mechanics of what have come to be called jammed packing. Their conjecture was as follows: of the great many possible jammed packings into which such a collective system might become trapped, in the vicinity of a jamming transition all packings become equally likely. The Edwards conjecture was validated only recently (*74*). In these inanimate inert systems, the place of energy, *E*, in thermal systems is then taken by local available volume, *V*. This assertion leads to the definition of an effective temperature, *T_eff_*, based upon the statistics of volume variation in jammed packings. Specifically, if in thermal systems,

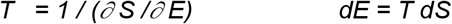

then in these granular athermal systems,

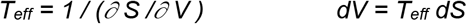

where *S* is the configurational entropy.

These notions of jammed packings, configurational entropy, and an effective temperature were subsequently extended to the living epithelial monolayer by Atia et al (*26*). They showed that just as volume variation follows a *k-gamma* distribution and maximizes configurational entropy in the jammed collective inanimate system(*27, 72*), so too does cell shape variation in the jammed confluent cellular system, both *in vitro* and *in vivo* (*26*).

###### *T_eff_* in tumor invasion dynamics

Using hard spheres in solution as a model system for jamming, one axis of the jamming phase diagram is typically given by *k_B_T/U* where *k_B_* is the Boltzmann constant, *T* is the thermodynamic temperature and *U* in interparticle attractive energy. This ratio is akin to an effective temperature in these systems. When temperature is higher or the particles are less attractive, the system tends to be less jammed and more fluid like. In our computational model system for tumor dynamics (Fig. 5), we find a similar balance between cellular propulsion and cell adhesion. If adhesion is kept constant as propulsion increases, we find that the cellular collective fluidizes. Indeed, MDA-MB-231 cells are both less adhesive (smaller effective interparticle energy) and more propulsive (higher effective temperature) than MCF-10A cells (*51*). In concert with that expectation, MDA-MB-231 cells fluidize in the same surrounding matrix more easily than do MCF-10A cells (Figs. 2 and 3).

###### Box 2: Effective Confinement Pressure

A variety of physical forces act to confine and direct collective cellular behavior. For example, cells within the tumor experience compressive stress due to uncontrolled growth. Indeed, Jain and colleagues have shown that the tumor interior develops compressive stresses large enough to collapse the intra-tumor vasculature (*22, 23, 75*). Literature developed by us (*16, 17, 20, 76*) and others (*31, 77*) shows that these compressive stresses can lead to systematic decreases of cell and nuclear volumes with increasing compressive stress. Additionally, within multicellular tumor models – and within the human tumor explant – nuclear volume varies appreciably and systematically both in space and time (*16*). As such, changes in cell and nuclear volume are sensitive to the local microenvironment (*20, 78, 79*), although whether or not they can serve as a remote pressure sensor, as some suggest (*31*), remains debatable. It is notable that, across these different systems, cell volume appears to change in close accord with the well-known Boyle van ‘t Hoff relationship (*20, 31, 76, 77*). These volume changes also appear to be associated with changing mechanical properties of the cell, with a strong increase in cell stiffness as cell volume decreases. From the Boyle van ‘t Hoff relationship, calculation of the bulk osmotic modulus, *B*, is straightforward (*76*):

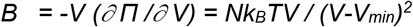

where *Π* is the osmotic pressure, *N* is the total number of osmolytes, *k_B_* is the Boltzmann constant, *T* is the temperature, *V* is the volume, and *V_min_* is the osmotically inactive volume.

For reasons that remain unclear, both cortical and cytoplasmic stiffness are orders of magnitude smaller but follow the same functional trend (*20, 76*). As is described below, increasing cell stiffness may influence jamming behavior and as such understanding these forces remains critical.

###### *P_conf_* in tumor invasion dynamics

In jamming behavior of colloids, micro-gels or many collective systems, both particle number density and particle stiffness play critical roles; higher number densities and stiffer particles tend to promote jamming (*80*).When these factors are held constant but collagen density is high, our computational model shows that the cellular collective can be either solid-like or fluid-like depending on propulsion (and therefore *T_eff_*; Box 1) but cell escape as a gas is not possible (Fig. 5). However, when collagen density is lowered, the cellular collective can become gas-like (depending on *T_eff_*), in which case cells escape readily. Changing collagen density in this computational model is akin to changing an effective confinement pressure, <u>*P_conf_*</u>, in which case the matrix is imagined to comprise a vessel that acts to confine the jammed collective. In concert with these notions, experiments show that MDA-MB-231 cells are softer and exert greater propulsive forces than do MCF-10A cells (*51*) and, as expected, escape more easily into an equivalent matrix (Figs. 3 and 4).

## General

The authors gratefully acknowledge Elizabeth Bartolak-Suki (Boston University) for expert assistance with spheroid histology, and Lauren O’Keeffe (Cornell University) for assistance with spheroid embedding and cell counting.

## Author contributions

W.K., J.F., J.J.F designed the study; J.F., Y.L.H, J.A.M., J.A.P, D.R. performed experiments; C.S. and Z.C. performed simulations; W.K., J.F., Y.S., S.K. analyzed data; W.K., J.F., A.F.P., J.P.B., J.J.F. interpreted data; W.K, J.F, A.F.P, J.J.F. wrote the manuscript; W.K., J.F., C.S., Y.L.H., J.P.B., M.G., Z.C., M.Z., A.F.P, J.J.F. reviewed and edited the manuscript. J.J.F oversaw the project.

## Data and materials availability

Data supporting the findings of this study is available from the corresponding author upon request. The computational agent-based model is available from GitHub at https://github.com/SpataC/2D-Cell-simulation.

## SUPPLEMENTARY METHODS

### Collagen Structure and Mechanics

Acellular collagen gels at concentrations of 1-4 mg/mL were prepared using the materials described in the main text and polymerized within PDMS molds to form cylindrical plugs with a diameter of 9mm and height of 3mm. After collagen self-assembly, the gels were kept hydrated via addition of 2 mL of 1x PBS. Structural and mechanical characterizations were carried out by using previously published methods (*69*), briefly summarized below.

#### Structure

The microstructural features of acellular collagen networks were imaged using a Bruker Ultima Investigator multiphoton microscope (MPM) and a 60x oil-immersion objective. Image stacks of dimensions 78.9 μm × 78.9 μm × 10 μm (x-y-z) were acquired with 1 μm steps at a resolution of 0.076 μm/pixel. Masks that identify collagen fibers from second harmonic generation (SHG) signal were generated using a global intensity threshold that was determined empirically (*69, 81*). Collagen gels were treated as biphasic mixtures (*71*) made of two constituents: a solid (*s*, collagen) and a fluid (*f*, 1x PBS). The mixture is fully saturated, that is *V^s^* + *V^f^* = *V* or *φ*^*s*^ + *φ*^*f*^ = 1, where *V^α^* and *V* represent constituent and total volumes, *φ^α^* = *V^α^*/*V* represent volume fractions with *α* indicating either solid (*s*, collagen) or fluid (*f*, 1x PBS) constituents. Due to the random orientation of self-assembled collagen fibers, the volume fraction of collagen can be calculated from the respective area fraction (*82*). Hence, we computed *φ^α^* = *V^α^*/*V* ≈ *A^α^*/*A*, here *A^s^* represents the masked area occupied by collagen fibers and *A* represents the total imaged area (78.9 × 78.9 μm^2^). The fluid volume fraction – or porosity – was then calculated as *φ^f^* = 1 − *φ^s^*. Individual collagen fibers were identified using CT-FIRE, a validated algorithm for segmentation of microscopy images and extraction of fiber geometry and alignment (*70*). Examples of collagen fiber identification via CT-FIRE segmentation are shown in Figure 4D. Collagen fiber density was calculated for each slice as the number of segmented fibers divided by the sampled volume (78.9 × 78.9 × 1 μm^3^) and averaged across each image stack.

#### Mechanics

The macroscopic properties of cylindrical collagen samples polymerized within PDMS wells were estimated using uniaxial confined compression testing (*83*) and continuum biphasic modeling (*84*). Experiments were carried out using a commercial DHR-2 rheometer (TA Instruments, New Castle, DE) equipped with a porous sintered steel mesh. Stress relaxation steps were imposed by applying a deformation equal to 3% of the unloaded gel height at a rate of 1%/s, followed by a 180 second hold. The axial force during each step of compression was recorded by the rheometer and six steps of compression were repeated for a final deformation of 18%. The bulk mechanical behavior measured under compression was modeled by separating two contributions: a solid stress generated by the collagenous matrix and a transient pressure generated by the fluid filling the interstitial space. Due to the experimentally observed nonlinear responses, the solid constituent was treated as hyperelastic, hence its mechanical response to deformation depends on a free energy density function *W*, which was modeled using the functional form originally proposed by Yeoh (*85*)

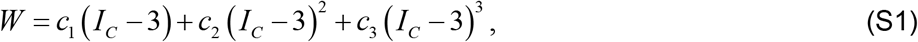

where *I_c_* = *tr***C** represents the first invariant of the Cauchy-Green tensor **C** = **F**^*T*^**F**, and **F** is the deformation gradient which, in confined compression, assumes the form **F** = diag(1,1,*λ*) where *λ* = *h*/*H* is the ratio between loaded (*h*) and unloaded (*H*) gel thickness – or axial stretch – imposed by the rheometer. The material parameters *c*_1_, *c*_2_, and *c*_3_ were subjected to the constraints *c*_1_, *c*_3_ > 0 and *c*_2_ < 0 (*85*). The fluid transport properties of collagen gels were instead described by a hydraulic permeability tensor **k**, which – according to Darcy’s law – regulates hydraulic flow in porous media in response to pressure gradients (*82*). Here, we assumed an isotropic and strain-independent hydraulic permeability, that is

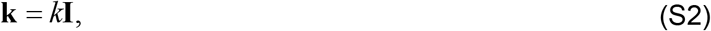

this assumption is due to the fact that collagen gels are highly porous materials made of randomly oriented fibers. Unknown model parameters were calculated by adopting a two-step fitting procedure of stress relaxation data (*71*): material parameters from the Yeoh model (*c*_1_, *c*_2_, *c*_3_) were determined by fitting steady-state data from the six stress relaxation steps, while the hydraulic permeability (*k*) was determined by fitting the corresponding transient responses. In addition to control gels at collagen concentrations between 1 and 4 mg/mL, the above methods were applied also to collagen gels that were chemically cross-linked using 0.2% glutaraldehyde (GA). Cross-linking does not change the microstructure and porosity of collagen gels (*86*), but allows for a more accurate determination of the hydraulic permeability by preventing plastic deformations upon compression (*69*). Further details on both experimental and modeling techniques can be found in (*69*).

### Agent-based modeling

We developed a computational model that represents a hybrid between vertex (*25*) and particle-based (*48*) approaches. Using a 2D computational domain, a cellular collective was generated via Voronoi tessellation and embedded within a collagen matrix, which is represented by randomly distributed collagen posts. The cell collective is modelled as a confluent tissue, with no birth and death events, and is considered as homogenous in terms of cell properties such as adhesion strength and inherent motility. By varying parameters such as cell motility and collagen fiber density, the model simulates various types of invasive behavior of a tumor cell cluster into surrounding ECM. The key features of the model are described below.

#### Cell mechanics

The cell membrane is modeled as viscoelastic, with neighboring points on the membrane interacting through both an elastic force 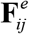 and a viscous force 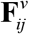 that act along the cell perimeter (*87*). These are respectively described by the following equations,

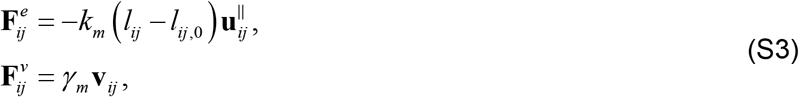

where *k_m_* is the elastic constant for the cell membrane, *l_ij_* is the current distance between adjacent points *i* and *j* on the membrane, *l*_*ij*,0_ is the respective equilibrium distance, and 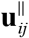 is the unit vector parallel to the segment *ij*, while *γ_m_* is the viscosity coefficient for the cell membrane, and **v**_*ij*_ is the relative velocity of point *j* with respect to point *i*. Moreover, a bending force 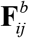 is introduced to correct for any inward or outward bending of the membrane during the simulation. This force is described as follows,

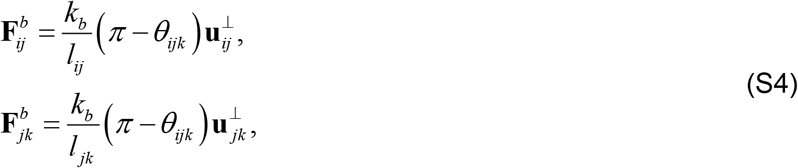

where *k_b_* is a bending stiffness of the cell membrane, *l_ij_* and *l_jk_* are the lengths of segments *ij* and *jk*, *θ_ijk_* is the angle between segments *ij* and *jk*, and 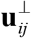 and 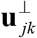 are unit vectors normal to segments *ij* and *jk* pointing towards the cell cytoplasm. It should be noted that these forces are aimed to align adjacent segments to achieve a smoother cell surface.

The cell body is also modeled as viscoelastic, with two main types of forces governing its mechanical behavior. First, an area-driven force 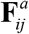 which represents the surface tension resisting to changes in the cell area. This is described as follows,

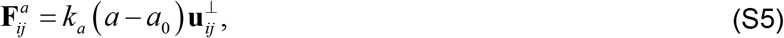

where *k_a_* is the elastic constant for the cell body, *a* is the current cell area, *a*_0_ is the equilibrium cell area, and 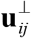 is the unit vector normal to segment *ij*. Second, an elastic force 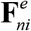 and a viscous force 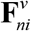 act along the actin fibers distributed radially along the cell cytoskeleton (*87, 88*). These fibers connect the nucleus (*n*) with a given point (*i*) on the cell membrane. These are respectively described by the following equations,

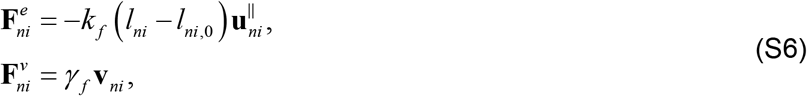

where *k_f_* is the elastic constant for actin fibers, *l_ni_* is the current fiber length, *l*_*ni*,0_ is the equilibrium fiber length, 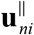 is the unit vector parallel to the segment *ni*, while *γ*_*f*_ is the viscosity coefficient for actin fibers, and **v**_*ni*_ is the relative velocity of point *i* on the membrane with respect to point *n*, the nucleus.

Each cell possesses an inherent or self-propelled motility that can be tuned via the propulsion parameter *v*_0_, similar to the one described by Bi et al.(*25*). Cell *n* receives a randomly generated polarity vector (cos*θ_n_*, sin*θ_n_*) along which the cell exerts a self-propulsion force with a constant magnitude *v*_0_ / μ, where μ represents the inverse of the frictional drag coefficient (*25*). The polarity vector undergoes rotational diffusion by a Gaussian white-noise process with mean 0 and variance *D_r_*, where *D_r_* is the rotational noise strength and is kept constant throughout the simulation.

#### Cell-cell interactions

Physical interactions between cells are modelled in the form of a Lennard-Jones potential (*89*). While the rejection component in the potential prevents cells from overlapping, the attraction component models cell-cell adhesive forces present in tissues. A notable example is given by the adherens junctions that are known to maintain structural stability of cell monolayers and allow functions such as cell communications and collective movements (*90*). The inter-cellular force **F**_*LJ*_ arising from the Lennard-Jones potential is described as follows,

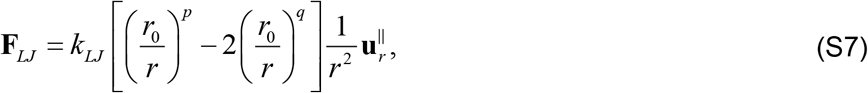

where *k_LJ_* is the constant that characterizes the strength of the interaction, *r* is the current separation distance (minimum distance between points on the membranes of two interacting cells), *r*_0_ is the maximum separation distance for rejection, 2*r*_0_ is the maximum distance for interaction, and 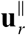 is the unit vector parallel to the separation distance. The exponents *p* and *q* were assigned values, respectively, of 6 and 3 that control the strength of cell-cell interactions.

#### Extracellular environment

The extracellular matrix (ECM) environment is assumed to be composed only of collagen posts, which are modeled as fixed elastic blocks that are randomly distributed around the cell cluster. Varying the density of collagen blocks allows one to vary the collagen density within the ECM. In this model, only physical interaction forces between cell and ECM are captured, and therefore does not consider active interactions such as the potential lysis of collagen by migrating cells. As such, cells have minimal interactive forces with the collagen blocks, consisting of small repulsion forces at the interface of cell surface-collagen surface, to ensure no spatial overlap. These interactions are modelled with a Lennard-Jones potential as described in Equation S7, with exponents *p* and *q* with values of 2 and 0 (no cell-ECM adhesion), respectively.

#### Model configuration

The initial configuration of the model is constituted by 152 cells arranged in a circular collective within an ECM environment represented by collagen blocks. Cell motility is represented by the propulsion vector *v*_0_, which is homogeneous in magnitude across the cell collective, and with *v*_0_ increasing as the cells comprising the collective become more and more motile. All the parameters described above are homogeneous across the cell population with values specified in the next section, the only major differentiator being the random motility orientation for each cell. Specifically, the initial model configuration is generated as follows. A number of points (n = 340) are seeded in a rectangular environment (261 × 160 pixels). Following a Poisson disk sampling to ensure a more uniform distribution of the generated points, a Voronoi tessellation is created based on these cell centers. The generated polygons are then shrunk isotropically to ensure a distance *ɛ* > 0 between all neighboring edges. The cell membrane is simulated by placing 25 equidistant points (indicated as *i*, *j*, *k* in the sections above) on each polygon’s sides. A disk of radius 73 pixels is used to sample the cells within a circular collective, leaving the remainder of the environment empty. This reduces the number of cells from the initial 340 in the whole environment to a total of 152 cells within the circular disk. The simulated ECM environment is obtained by seeding a specific number of points (depending on the desired collagen density) in an extended environment (520 × 320 pixels) and generating spherical collagen blocks, with a fixed radius (7 pixels) and a fixed number of exterior points (12 points). Starting from this initial configuration, the model simulates the invasive behavior of the cell collective over time.

#### Simulation set-up

The simulation is advanced by adopting a semi-Implicit Euler scheme:

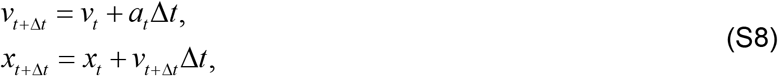

At time *t* + Δ*t*, the velocity of any point on the cell membrane *v*_*t*+Δ*t*_ is computed using the previous timepoint’s velocity *v_t_* and acceleration *a_t_*. Acceleration for each cell is the total force applied on the cell at time *t* divided by the mass. For simplicity, the mass was taken to be 1 for all cells. Following the computation of the updated velocity, the next position (*x*_*t*+Δ*t*_) of each point on the cell membrane is computed to advance the simulation. The whole system is damped by a parameter *v_decay_* that serves as a correction for the acceleration generated by errors in the advancing method. This damping correction is implemented by updating *v*_*t*+Δ*t*_ (S8-1) with 1−*v_decay_* at each time step. Based on preliminary simulations, here we used *k_m_* = 0.25, *l*_*ij*,0_ = 1.48, *γ_m_* = 5, *k_b_* = 2×10^−3^, *k_a_* = 1×10^−3^, *a*_0_ = 10^2^, *k_f_* = 5×10^−4^, *γ_f_* = 0.58, *k_LJ_* = 0.75, *r*_0_ = 1, *v_decay_* = 10^−4^, all expressed in arbitrary units (A.U.).

#### Phase diagram simulation and robustness testing

To investigate the effect of varying cell motility and collagen density on spheroid invasive behaviors, we varied the cell propulsion *v*_0_ in the range [0.6, 1.18], with a step increment of 0.02 to simulate the effect of increased cell motility. Meanwhile, the effect from the surrounding ECM that the cell collective experiences is tuned by modifying the spatial density of collagen blocks. For each combination of cell motility and collagen density, the 95^th^ percentile of radial cell positions (**p**^95th^) at the end of each simulation was computed and the cumulative distribution of **p**^95th^ resulting from all simulations (Figure 5C) was used as a metric to distinguish between material phases (solid, liquid or gaseous) exhibited by the invading cell front. By using **p**^95th^, we chose to examine only the position of the 5% outermost cells as the minimal fraction of the cell collective that is needed to capture the periphery of the tumor near the cell-ECM interface. In addition, **p**^95th^ provides the most straightforward way to identify what type of phase transitions occurred in the cell collective. Based on the resulting distribution, the thresholds for each of the phases were set as the 34^th^ percentile for the liquid phase, and as the 64^th^ percentile for the gaseous phase (Figure 5C). All simulations were run in 10 replicates, each using different sets of collagen post distribution and cell motility orientation. Each data point from the phase diagram in Figure 5D represent the averaged result from these 10 configurations.

To assess the effect of the arbitrary thresholds chosen based on the radial distance from the center of the cell collective on the identification of distinct material phases, we performed two separate tests. First, we performed a sensitivity analysis aimed at assessing the influence of different threshold values on the phase diagram. We found that the phase diagram is stable, and not sensitive to a number of threshold changes (Table S2). Second, we assessed whether a criterion based solely on radial distance reflects differences in cell shape, thereby separating different material phases. Based on a fixed initial configuration of both cells and collagen, we ran a new set of simulations in correspondence of all combinations of cell motility and collagen density. For such combinations, which reflect all the points on the phase diagram, we assessed the material phase based on the radial distance criterion and calculated the average cell shape index. This average cell shape, defined as 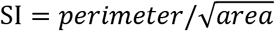 for cells in 2D, is calculated for cells within a 95% radial distance from the center of each collective, and allowed characterization of the bulk of the collective that is not fully captured using the radial distance criterion. By comparing the mean SI of cell collectives that are categorized as solid-like or liquid-like by the radial distance criterion, we found that liquid-like cell collectives are on average more elongated and have more variables shapes (SI^liquid^ = 4.75 ± 0.35) with respect to their solid-like counterparts (SI^solid^ = 4.50 ± 0.11, p = 1.95×10^−6^, H0: SI^liquid^ = SI^solid^), as expected from modeling studies (*25*). It should be noted that a mean SI is not computed for individual gas-like cells, as it is defined only for multicellular collectives in which individual cells are fully surrounded by neighboring cells.

## SUPPLEMENTARY FIGURES

**Fig. S1.**
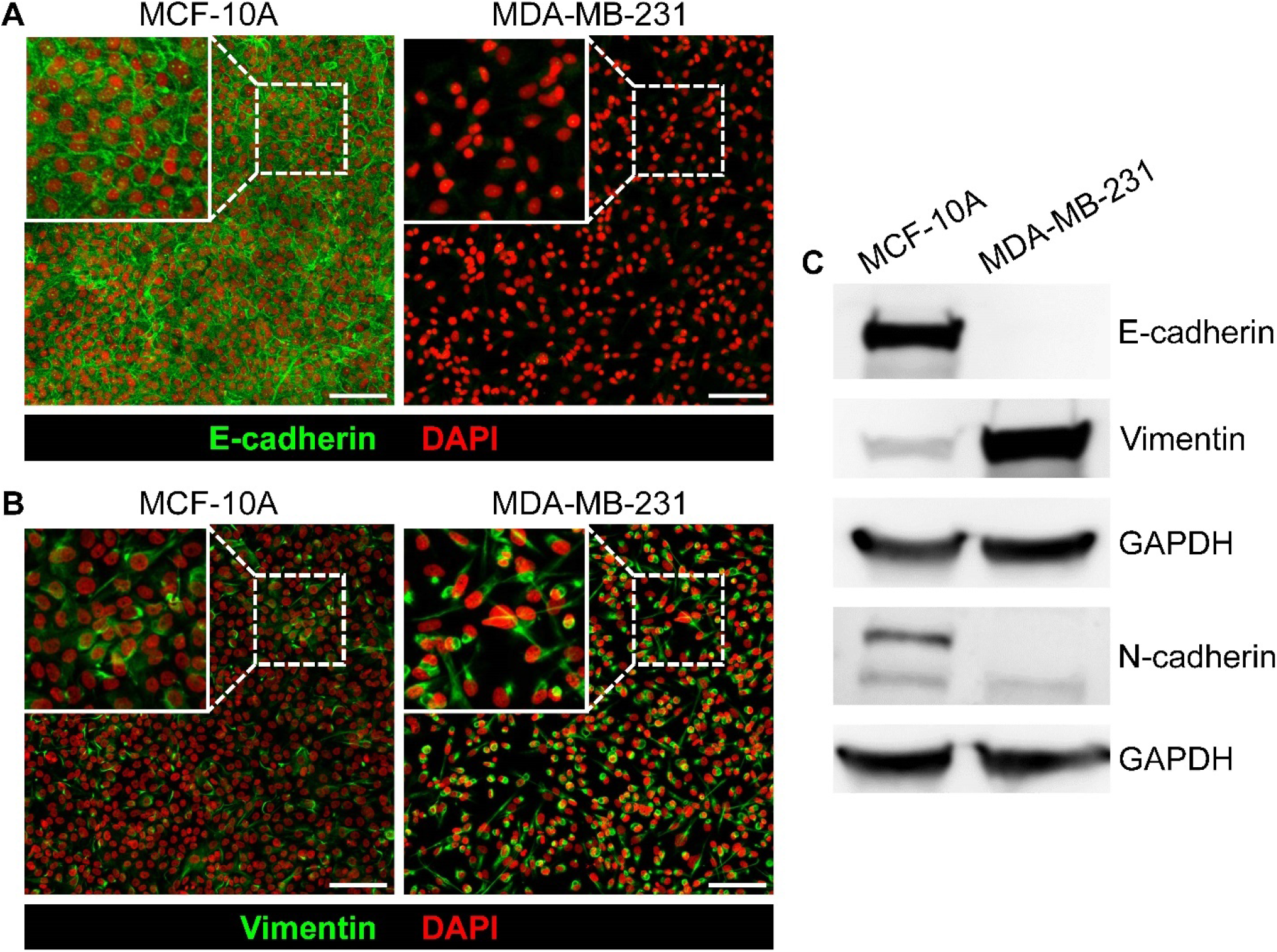
Characterization of MCF-10A and MDA-MB-231 cell lines. Representative immunofluorescence (IF) staining for (**A**) E-cadherin and (**B**) Vimentin shows that MCF-10A cells display predominantly epithelial characteristics while MDA-MB-231 cells show predominantly mesenchymal characteristics. Scale bars represent 100 μm. (**C**) Western blot analysis of MCF-10A and MDA-MB-231 monolayers confirms IF findings. E-cadherin is expressed only in MCF-10A but not MDA-MB-231 cells, while vimentin is expressed at a low but detectable level in MCF-10A cells and at a high level in MDA-MB-231 cells. Both cell types show expression of N-cadherin, with higher expression in MCF-10A cells. Together these data indicate that MDA-MB-231 cells are fully mesenchymal, while MCF-10A cells are mostly epithelial but with some aspects of an early partial EMT phenotype, as is typical for this model epithelial cell line.

**Fig. S2.**
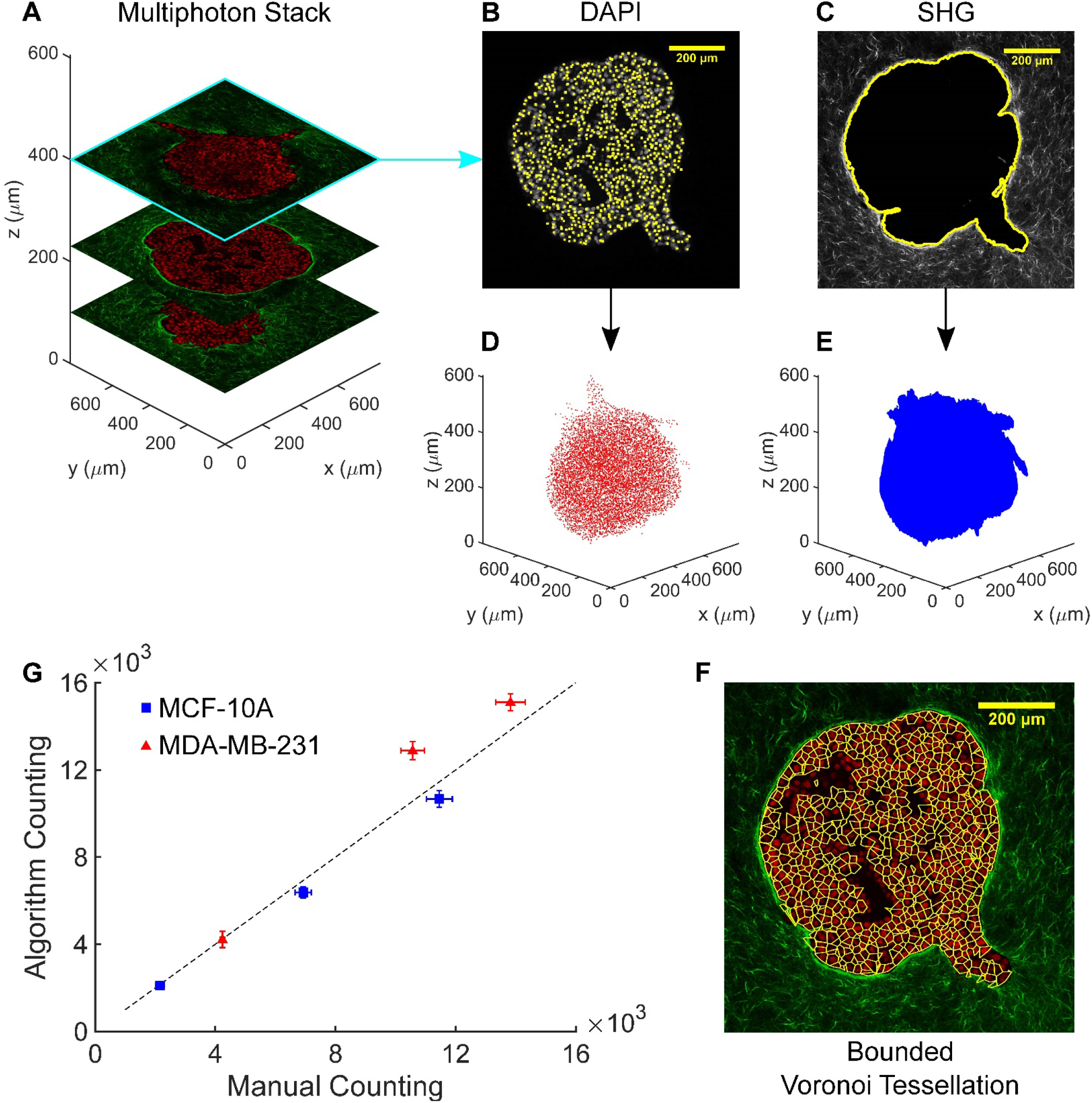
Workflow for 3D cell identification and cell shape estimation in multicellular spheroids. Here, the workflow is illustrated for MCF-10A macro-spheroids in collagen, but was also applied to MCF-10A microspheroids and MDA-MB-231 macro-spheroids alike. (**A**) A representative multiphoton stack of a MCF-10A macro-spheroid cultured in 2 mg/ml collagen is split into two channels, carrying separate signals: (**B**) fluorescence from DAPI-stained nuclei and (**C**) second harmonic generation (SHG) from collagen. In the case of MCF-10A micro-spheroids in Matrigel and Alginate, the two channels are given by fluorescent and bright-field images (cf. Figure 1). The two channels are processed, respectively, to identify (**D**) the 3D position of cell nuclei and (**E**) the 3D spheroid boundary using a custom algorithm. Note that the representative slices (**B-C**) displaying 2D projections of identified nuclei centers and spheroid boundary are shown overlapped onto corresponding multiphoton images. (**F**) Data on nuclei location and spheroid boundary are combined to generate a bounded Voronoi tessellation that partitions the spheroid into individual cells, from which cell volume and shape metrics are derived. Here, the 2D cross-section view of resultant Voronoi cells are shown superimposed on the corresponding multi-photon image. We note that empty spaces within the spheroid (cf. Fig. S6) as well as its neighboring cells are excluded from further analysis to minimize local overestimation of cell shape (**G**) Our custom 3D nuclei detection algorithm was validated by comparing the number of cell nuclei counted by the algorithm in fixed and cleared spheroids with the number of cell nuclei estimated for spheroids that were dissociated via trypsination and manually counted using a hemacytometer. Over a wide range of spheroid sizes, using both the MCF-10A (blue squares) and MDA-MB-231 (red triangles) cell lines, manual and algorithm counts lie close to the identity line (dashed black line). Spheroids used for counting validation were formed starting from a variety of cell numbers (from ~10^3^ to ~10^4^) at the time of seeding which increased (to ~2×10^3^ to ~1.5×10^4^) over the course of the 48 hours during which spheroid formation occurred (cf. Fig. S6). Both cell couting methods used spheroids from the same batch and cells at the same passage. Data are presented as mean ± SEM (n = 6-10 for manually counted spheroids and n = 3-4 for automaticaly counted spheroids of all sizes).

**Fig. S3.**
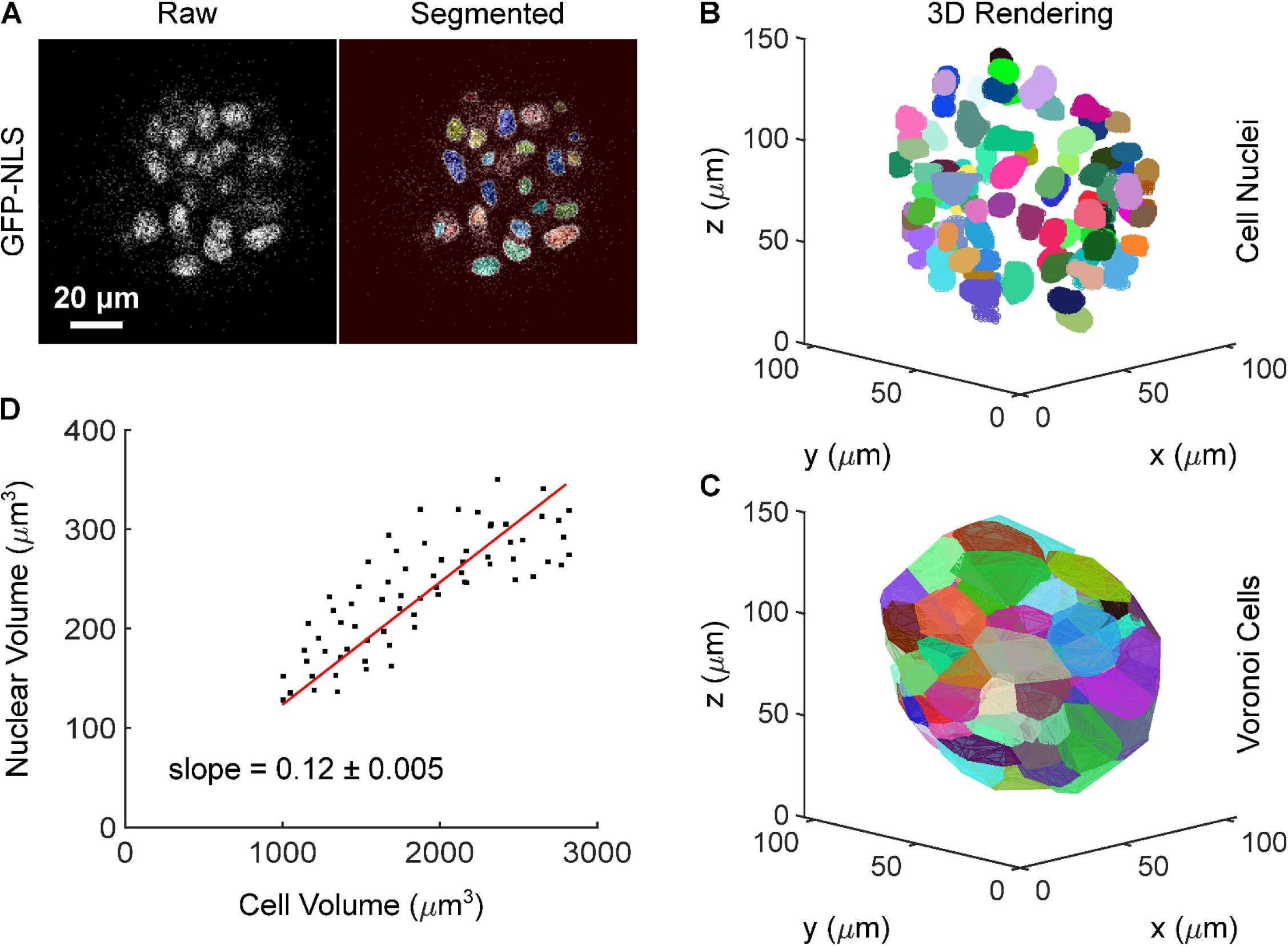
Cell volume from 3D Voronoi tessellation tracks nuclear volume in multicellular spheroids. (**A**) Equatorial cross-section of a representative confocal microscopy image from an early GFP-NLS labelled MCF-10A micro-spheroid juxtaposed to the segmented image, in which each identified nucleus is labeled with a different color. (**B**) 3D rendering of an early MCF-10A spheroid in which cell nuclei are reconstructed from segmentation of confocal stacks. (**C**) 3D rendering of the same spheroid in which Voronoi cells are obtained using the workflow outlined in Fig. S2. (**D**) Voronoi cell volume plotted against nuclear volume for individual cells show that the two measures are linearly related (R^2^=0.95).

**Fig. S4.**
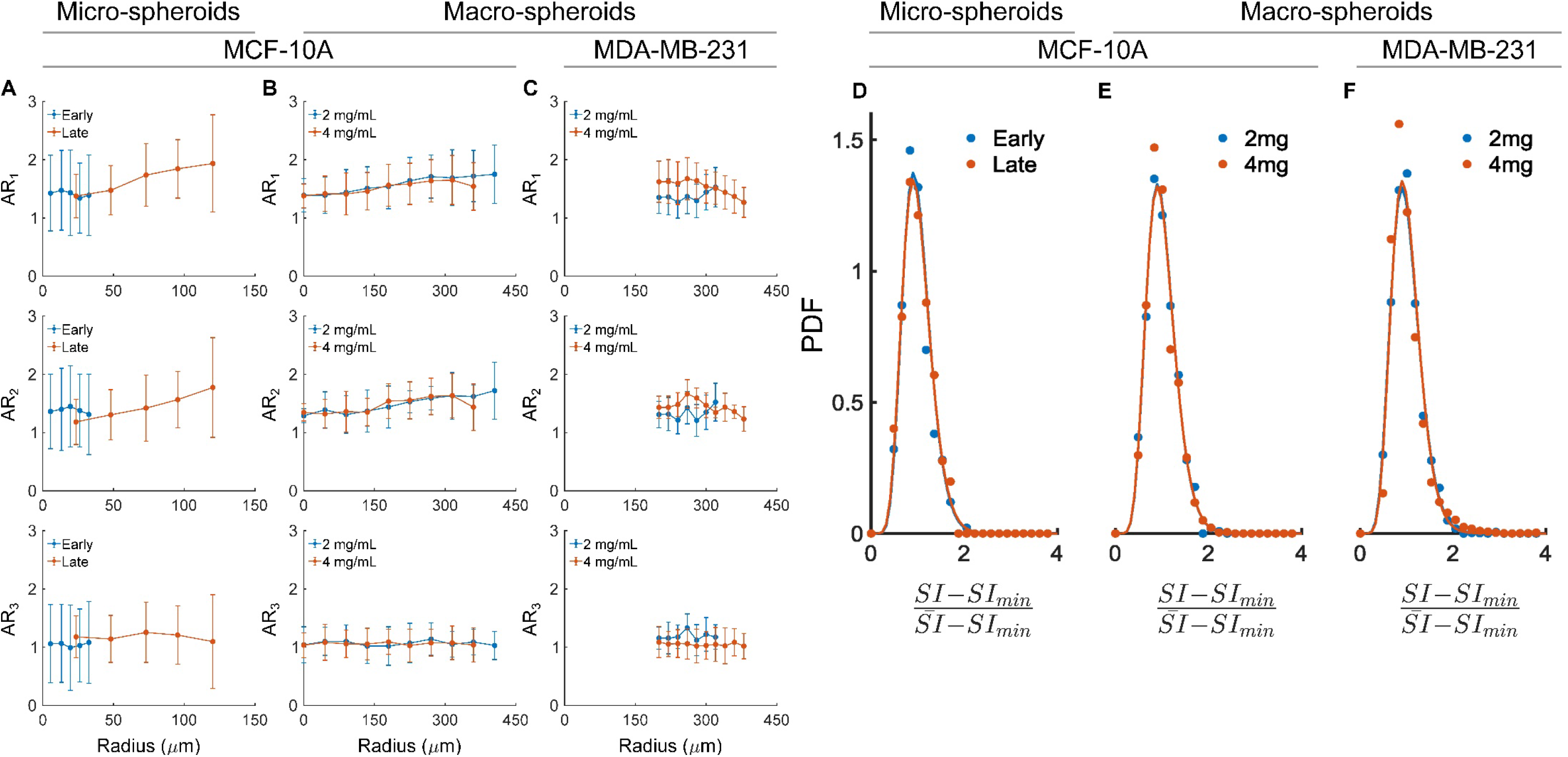
Aspect Ratio (AR) and Shape Index (SI) distributions within multicellular spheroids. Following Atia et al.(*26*), we used tesselletad Voronoi cells to quantify cell shape in terms of AR, which is closely associated with the second moments of inertia **I** and can be described using the diagonal terms of the second-order tensor **I**= *diag* {*I*_1_,*I*_2_,*I*_3_}. We treated each cell as a polyhedron of uniform density, and calculated respective moment of inertia tensors acoording to Mirtich et al.(*91*). The three AR metrics, arranged in descending order, are AR_1_=*I_1_*/*I_2_*, AR_2_=*I_1_*/*I_3_* and AR_3_=*I_2_*/_3_. In addition to serving as a structural signature of unjamming, respective change in the three ARs also inform about nature and orientation of cell shape changes. (**A**) Upon MCF-10A micro-spheroid growth from early to late stage, both AR_1_ and AR_2_ show development of increasing radial gradients towards larger AR for cells in the periphery of late stage spheroids. The increase in AR are consistent with increase in cell shape index (cf. Figure 1), and support the existence of more elongated cells – that is, unjammed – peripherial cells. In addition, the radial increase in both AR_1_ and AR_2_ highlights the existence of more rounded cells in the spheroid center (AR near 1) and more rod-like cells at the spheroid periphery, in line with interpretation of cell polarization outward to invade into the matrix. (**B**) Similar, but less pronounced, gradients can be seen in MCF-10A macro-spheroids cultured in collagen, where AR_1_ and AR_2_ increase towards the spheroid periphery only for speroids embedded in 2mg/ml collagen, that is, for spheroids that undergo local unjamming with invasion. (**C**) Instead, a small radial decrease in AR_1_ is seen for cells from MDA-MB-231 macro-speroids in 4mg/ml collagen, which suggests a transtion to more rounded cell shapes in correspondece of the invasive protrusions, thus supporting the notion of a confinement-induced jamming transition. For each Voronoi cell, we computed the SI as *surface*/*volume>*^2/3^ and fitted SI distributions to the *k-gamma distribution* using maximum likelihood estimation [2]. Each distribution is generated by pooling data from all experiments and dividing it into 50 bins. Here we present the experimentally obtained PDFs as data points (circles) and show their respective best-fitting k-gamma distributions (solid lines of the corresponding color). (**D**) MCF-10A micro-spheroids shape distributions at the early and late stage fit for a k-value of 10.06 ± 0.05 and 10.01 ± 0.07, respectively (p=0.32). (**E**) MCF-10A macro-spheroids shape distributions in 2 and and 4 mg/ml collagen fit for a k-value of 10.14 ± 0.17 and 10.20 ± 0.11, respectively (p=0.51). **(F)** MDA-MB-231 macro-spehoids shape distributions in 2 and 4mg/ml collagen fit for a k-value of 10.38 ± 0.10 and 10.26 ± 0.15, respectively (p=0.24). Across cell types, experimental preparations, and invasion phenotypes, the distribution of cell shapes robustly followed a k-gamma distribution. Surprisingly, there was no statistically significant difference between the k-value that describe these distributions (H0: no difference in k=values between respective conditions). Average k-value across cell types and spheroid preparations is 10.19 ± 0.1, consistent with the k-value range from the theory of 3D inert matter packings from Aste and Di Matteo [3], and suggests that variations in individual cell shapes are perhaps due to short ranged interaction between immediate neighbors.

**Fig. S5.**
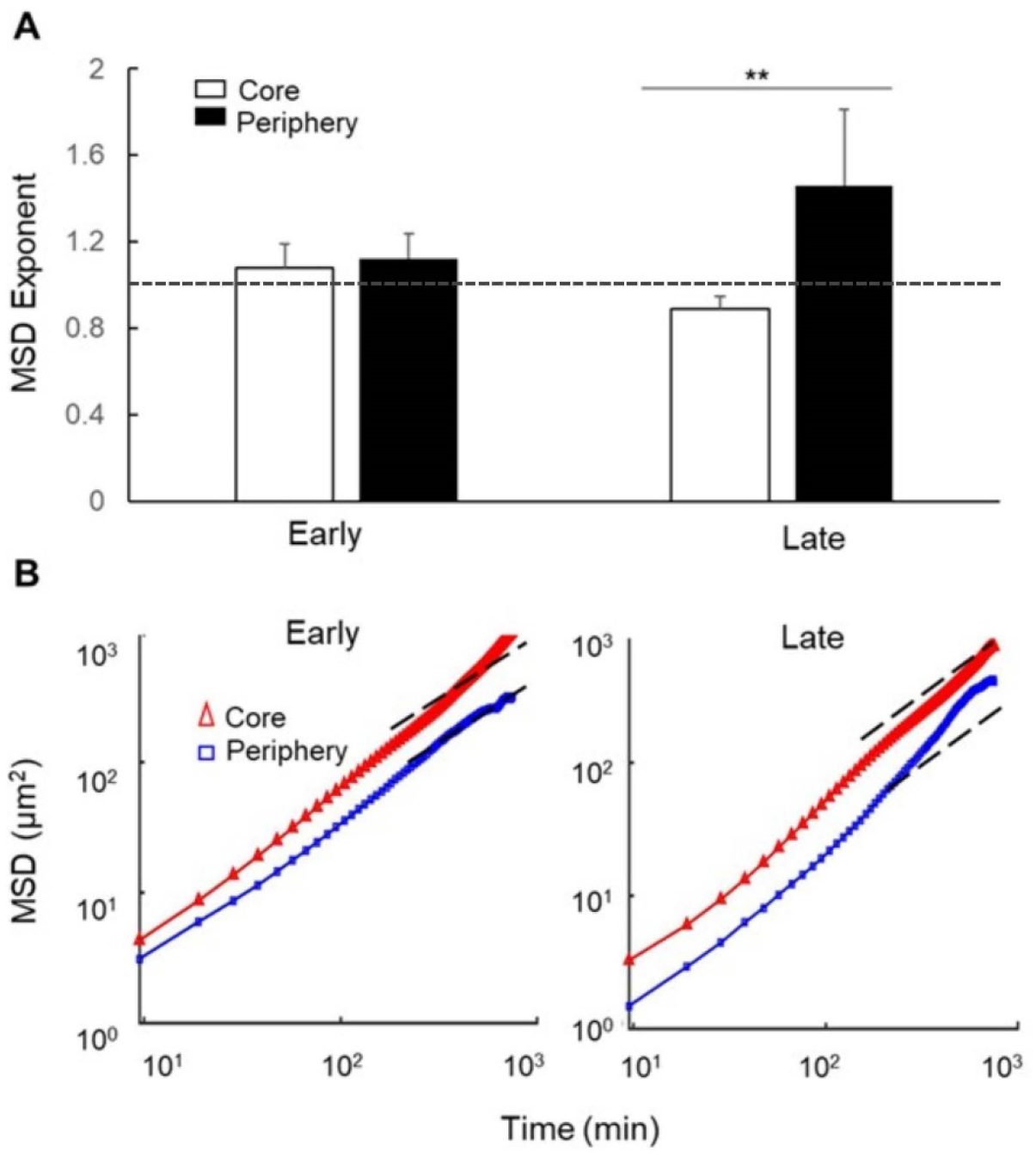
Migratory dynamics shows spatio-temporal dependence of cellular motion during micro-spheroid growth and invasion. (**A**) Exponent of mean squared displacements (MSD) show cellular motions are diffusive and homogenous in the early stage micro-spheroid (p=0.67), while spatial dependence in cell motion is clear in the late stage micro-spheroid (p<0.01). In the late stage micro-spheroid, cellullar motion is sub-diffusive (p=0.051, H0: exponent not different from 1) in the spheroid core suggestive of caging, while periphery cells are clearly super-diffusive. MSD exponent = 1 is indicated with a dashed line to guide the eye. MSD exponents are obtained from linear fitting of MSD as a function of time on a log-log plot. Core regions are defined as radius < 50% of initial micro-spheroid radius, and those above as periphery region. Data are presented as mean ± standard deviations. (**B**) Time and ensemble averaged MSD of cell trajectories from all spheroids over the observation window. The black dashed line show a slope of 1 (MSD exponent = 1, suggestive of diffusive motion) to guide the eye. (n = 5 for micro-spheroids at early and late stage respectively).

**Fig. S6.**
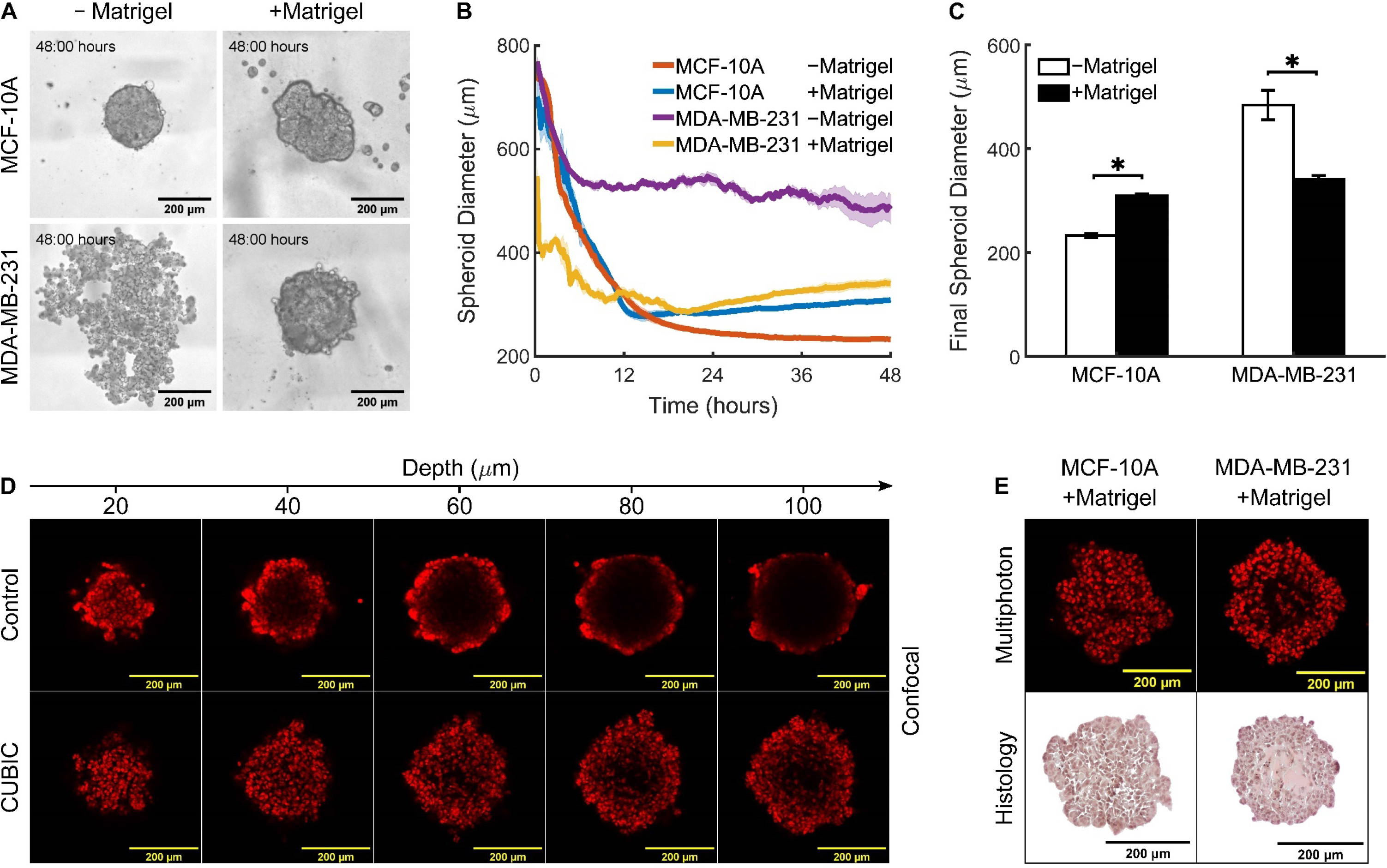
Formation and 3D imaging of MCF-10A and MDA-MB-231 macro-spheroids. (**A**) Representative DIC images of macro-spheroids formed in low attachment conditions for 48 hours starting from approximately 10^3^ cells in presence (+) or absence (−) of 2.5% Matrigel diluted in cell culture media. The presence of Matrigel leads to MCF-10A macro-spheroids that are larger and less regular in shape with respect to Matrigel-free controls. On the other hand, MDA-MB-231 cells form only loose aggregates in absence of Matrigel while compacting effectively in its presence. (**B**) Time-course of spheroid compaction over the course of 48 hours. The equivalent spheroid diameter *d* is calculated from the total cell area *A* thresholded from DIC time-lapse movies as 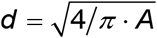. (**C)** Spheroid diameter at 48 hours of compaction quantifies the differences caused by the presence of Matrigel in the different cell lines. Both cell lines form spheroids of similar sizes in the presence of Matrigel (black bars). Data are presented as mean ± SEM (n = 3 for all groups). * indicates statistical significance at p < 0.05. (**D**) 3D scanning confocal microcopy images of representative MDA-MB-231 spheroids after fixation (Control) or fixation followed by optical clearing (CUBIC), shows that DAPI-stained nuclei (red) in the spheroid core are clearly visible only after optical clearing. The depth of the confocal stacks represents the distance from the spheroid surface. (**E**) Representative equatorial cross-sections of MCF-10A and MDA-MB-231 spheroids generated in presence of Matrigel show the distribution of cell nuclei using two methods: multiphoton imaging of DAPI-stained cell nuclei after CUBIC clearing (top), and van Gieson’s stained histology slides (bottom). Note that the hollow core of MDA-MB-231 spheroids is not filled by cells (identified by the dark nuclei) but rather is occupied by matricellular proteins (shown in red).

**Fig. S7.**
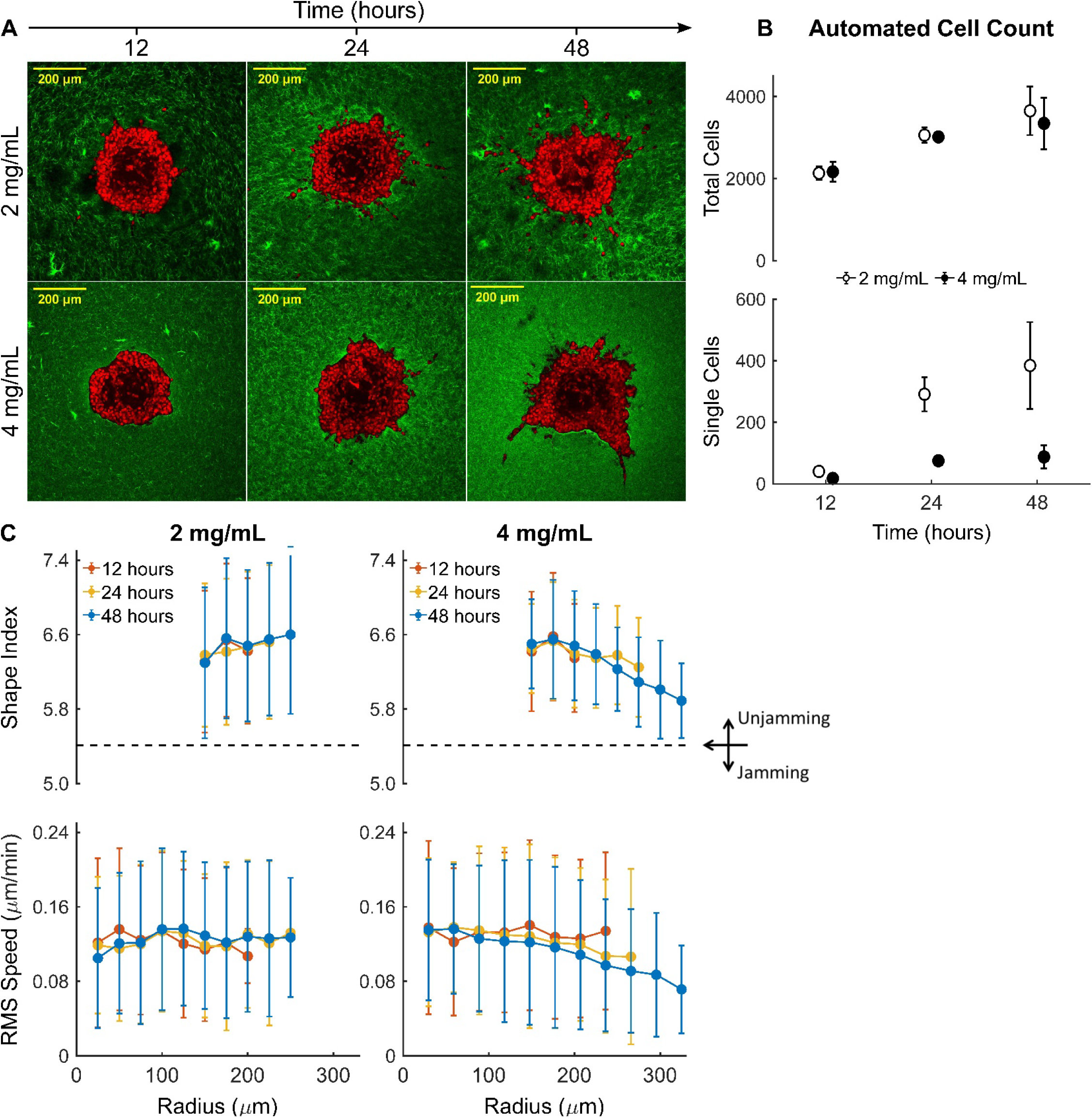
Over time, MDA-MB-231 macro-spheroids in high density collagen experience confinement-induced jamming at the invasive front. We carried out a time-lapse experiment, including continuous DIC imaging as well as fixation and optical clearing for multiphoton microscopy, for MDA-MB-231 spheroids at distinct time points. (**A**) Representative equatorial cross-sections of multiphoton images display DAPI-stained cell nuclei (red) and collagen fibers from SHG (green) from MDA-MB-231 spheroids embedded in 2 and 4mg/ml collagen and imaged after 12, 24, and 48 hours. (**B**) Total and single cell counts over time show that cell proliferation is quite similar between spheroids embedded in various collagen densities, while there is a significantly higher single cell escape over time in spheroids embedded in 2 mg/ml collagen. (**C**) Temporal evolution of cell shape (top) and migratory speed (bottom) in spheroids embedded in 2 mg/ml (left), and 4 mg/ml (right) collagen. In 2 mg/ml collagen, MDA-MB-231 cells that remain within the primary spheroid maintained homogenous radial distributions for both cell shape and RMS speed.In contrast, in 4 mg/ml collagen, MDA-MB-231 cells within the primary spheroid have similar homogenous distributions until 12 hours of culture, but progressively develop radially decreasing trends for both cell shape and RMS speed at 24 and 48 hours.The critical cell shape (SI = 5.4) for solid-fluid transitions is indicated by a horizontal dashed line, with increasing values indicative of unjamming and decreasing values indicative of jamming. Overall, we observe that ECM confinemnt induces a jamming transition, or confinement-induced jamming, at the invasive front.

**Fig. S8.**
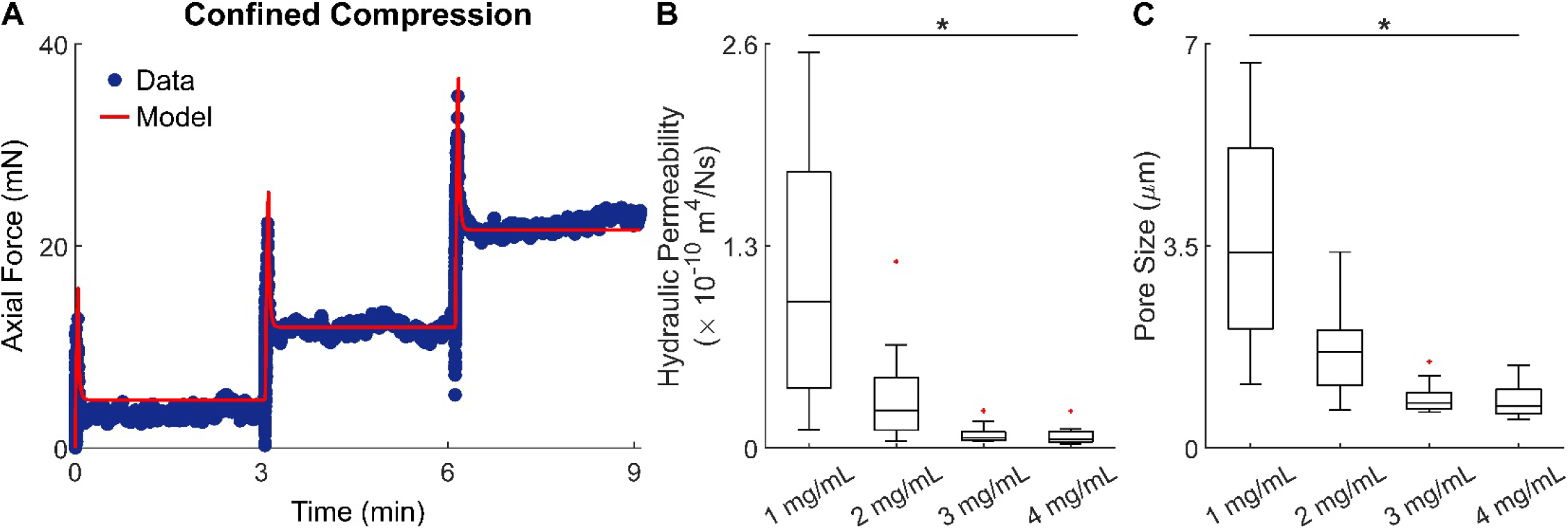
Increasing collagen density associates with a sharp decrease in hydraulic permeability and pore size. We performed structural and mechanical characterization of acellular collagen gels to show that microstructural, rather than mechanical, changes with increasing collagen density underlie the observed changes in MDA-MB-231 invasive phenotype. (**A**) Representative confined compression data from a 3 mg/ml collagen gel chemically cross-linked with 0.2% glutaraldehyde (GA). We recently showed that GA cross-linking allows for improved estimation of fluid transport properties in porous gels via continuum biphasic modeling [4]. (**B**) Box-Whisker plots of hydraulic permeability and (**C**) pore size show a consistently decreasing trend with increasing collagen concentration, which mirrors the microstructural – rather than mechanical – trends reported in Figure 4. Instead of measuring directly the pore size, we calculated it by using the Carman-Kozeny model which relates permeability, porosity, and pore size [5]. It should be noted that Sauer et al. [6] recently showed that GA cross-linking does not affect the pore size of collagen networks and that our calculated values are remarkably similar to their measure values. Permeability and pore size data are shown from 1 mg/ml (n = 9), 2 mg/ml (n = 9), 3 mg/ml (n = 10), and 4 mg/ml (n = 9) collagen gels. * indicates statistical significance at p < 0.05.

## SUPPLEMENTARY TABLES

**Table S1:** Increasing collagen density does not impact cell proliferation or spheroid growth rate. Both types of macro-spheroids (MCF-10A and MDA-MB-231) in both low and high collagen densities (2 and 4 mg/mL) were formed starting from similar cell numbers at initial seeding (Methods). Following 48 hours of spheroid formation and additional 48 hours of invasion in collagen, spheroids were fixed, optically cleared and imaged. Cell numbers within each macro-spheroid were counted using our custom cell identification algorithm (cf. Methods, Fig. S2). Spheroid radius was calculated from segmentation of DIC time-lapse movies to obtain the main spheroid area over time as 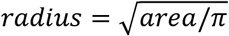. Radial growth rate is estimated from linear fitting of the spheroid radius over time. Increasing collagen density does not impact either cell count or radial growth rate while it affects the mode of migration (cf. Figures 2 and 3).

| Cell type | Collagen concentration | Cell count | p-value | Radial growth rate (μm/hr) | p-value | n |
| --- | --- | --- | --- | --- | --- | --- |
| MCF-10A | 2 mg/ml | 10369 ± 509 | 0.51 | 1.22 ± 0.16 | 0.24 | 3 |
|  | 4 mg/ml | 10679 ± 536 |  | 1.16 ± 0.12 |  | 3 |
| MDA-MB-231 | 2 mg/ml | 6230 ± 793 | 0.19 | 0.94 ± 0.27 | 0.53 | 3 |
|  | 4 mg/ml | 5536 ± 331 |  | 1.06 ± 0.15 |  | 3 |

**Table S2.** A sensitivity analysis shows that phase diagram is not sensitive to the threshold values used to define the different material phase states. Phase states of spheroids were determined from thresholding on distribution of simulated distance of the outermost 5% cells in the spheroid. Distances below the 34^th^ percentile, between the 34^th^ and 64^th^ percentile, and above the 64^th^ percentile lead to classification of migrating cells, repectively, as solid-like, fluid-like, and gas-like. We performed a sensitivity analysis to assess the dependence of the resulting phase diagram on the threshold values used to define the phase states, by applying ±2.5% changes to such thresholds. Results were quantified as percent change (%) in the number of points that have switched phase states (for example: solid to liquid, liquid to gas, etc.) from the total number of points in the phase diagram, due to the different threshold combinations. Our sensitivity analysis shows that the phase diagram is not significantly impacted by the exact values of threshold used to define the material states, with less than 5% of points changing material phase in correspondence of the highest change in threshold values tested.

| Gas Phase Threshold (%) | Liquid Phase Threshold (%) |  |  |  |  |  |
| --- | --- | --- | --- | --- | --- | --- |
|  | 31 | 32 | 33 | 34 | 35 | 36 |
| 62 | 4.872% | 3.590% | 2.821% | 2.051% | 2.051% | 3.333% |
| 63 | 3.077% | 3.077% | 2.051% | 1.026% | 1.026% | 2.308% |
| 64 | 3.077% | 2.051% | 1.026% |  | 0.000% | 1.795% |
| 65 | 4.872% | 2.821% | 1.795% | 0.769% | 0.769% | 2.564% |
| 66 | 4.872% | 3.846% | 2.821% | 1.795% | 1.795% | 3.590% |

## SUPPLEMENTARY MOVIES

**Movie S1:** Representative DIC time-lapse movies capture the migratory dynamics exhibited over the course of 48 hours by MCF-10A and MDA-MB-231 spheroids invading in 2 and 4 mg/ml collagen.

**Movie S2:** Representative simulation showing a cell collective exhibiting a solid-like non-invasive phase. Although cells remain migratory within the collective, the cell collective does not invade into the surrounding ECM. Cells are shown as orange particles, while green particles represent the surrounding ECM. This simulation corresponds to high collagen density (0.82) and low cell motility (0.2).

**Movie S3:** Representative simulation showing a cell collective exhibiting a liquid-like invasive phase. Cells slowly form streams that invade collectively into the surrounding ECM. Cells are shown as orange particles, while green particles represent the surrounding ECM. This simulation corresponds to high collagen density (0.82) and high cell motility (1.0).

**Movie S4:** Representative simulation showing a cell collective exhibiting a gas-like invasive phase. Cells from the collective invade into the ECM either as single cells or as small cell clusters. Cells are shown as orange particles, while green particles represent the surrounding ECM. This simulation corresponds to low collagen density (0.21) and high cell motility (1.0).

**Movie S5:** DIC time-lapse movies recorded over the course of 48 hours for MCF-10A spheroids embedded in graded concentrations of collagen (1 to 4 mg/ml). The periphery of MCF-10A spheroid remained solid-like and non-invasive in 4mg/ml, formed collective protrusions as collagen concentration was progressively reduced to 3 and 2mg/ml. Notably, as collagen concentration was further reduced to 1mg/ml, single cell detachment was observed while the main spheroid remained solid-like.

